# Pre-assembly of biomolecular condensate seeds drives RSV replication

**DOI:** 10.1101/2025.03.26.645422

**Authors:** Dhanushika Ratnayake, Marie Galloux, Sanne Boersma, Christina Sizun, Julien Sourimant, Anke J. Lakerveld, Matthijs J. D. Baars, Rupa Banerjee, Marko Noerenberg, Birgit Dreier, Sven Furler, Alfredo Castello, Andreas Plückthun, Jean-François Éléouët, Puck B. van Kasteren, Marie-Anne Rameix-Welti, Marvin E. Tanenbaum

## Abstract

During infection many RNA viruses, including respiratory syncytial virus (RSV), form specialized biomolecular condensates, inclusion bodies (IBs), where viral transcription and replication occur^1–4^. Paradoxically, high protein concentrations are typically required for condensate nucleation^5^, yet attaining sufficient protein levels in infection is thought to require IBs for viral transcription and replication. To uncover how viruses solve this paradox to establish IBs, we visualized early infection of RSV in real-time with single genomic viral ribonucleoprotein (vRNP) resolution. Our results reveal that IBs are nucleated from infecting vRNPs rather than *de novo* in the cytoplasm. IB nucleation further requires in-virion pre-assembly of viral protein-protein interaction networks on vRNPs to form ‘pre-replication centers’ (PRCs). PRCs are potent condensate nucleation seeds due to their resistance to disassembly and efficient recruitment of newly-synthesized viral proteins. The high protein affinity of PRCs also results in increased polymerase complex association, allowing efficient viral transcription even in the absence of IBs. Together, these activities create a feed-forward loop that drives rapid IB formation. Intriguingly, PRC assembly depends on in-virion viral protein levels and is highly heterogeneous among virions, explaining cell-to-cell heterogeneity in infection progression, and identifying heterogeneous virions as the origin of infection heterogeneity. Together, our results show that in-virion pre-assembly of PRCs kick-starts viral condensate nucleation upon host-cell entry, and explains cell-to-cell heterogeneity in RSV infection.

## Main

Viruses of the *Mononegavirales* (*MNV*) order, which include Ebola virus, measles virus (MeV) and respiratory syncytial virus (RSV), are among the most infectious human and animal pathogens. While effective vaccines and antiviral therapies are available for a number of viruses from this order, many remain untreatable and cause a major health and economic burden on society. Like other viruses of the *MNV* order, RSV has a non-segmented negative-sense RNA genome, encapsidated by viral nucleoprotein (N), together called the nucleocapsid (NC)^6–8^. In addition to N, the viral genome also associates with the viral RNA-dependent RNA polymerase (RdRp) complex, composed of the viral polymerase, ‘Large’ protein (L) and its essential co-factors, phosphoprotein (P) and the viral transcription factor M2-1. Together with the NC, these proteins form the viral ribonucleoprotein (vRNP) complex, the minimal infectious unit of RSV that functions as the viral transcriptase and replicase^9,10^.

Many viruses, including RSV, form cytoplasmic membrane-less compartments called viral factories or inclusion bodies (IBs), which are thought to act as the primary sites of viral transcription and replication^1–4^. While the importance of IBs for viral infection has been well documented, it is poorly understood how these organelles are initially nucleated. IBs are biomolecular condensates thought to form via liquid-liquid phase separation (LLPS) of the viral proteins, N and P^1,11,12^. However, phase separation reactions are very sensitive to the concentration of their constituent biomolecules; nucleation of condensates occurs when the concentration of the constituent biomolecules exceeds a threshold concentration, while condensates dissolve when the concentration drops below this critical concentration^13^. During early infection viral protein levels are very low, raising the question of how IBs can be nucleated. Moreover, IBs themselves are thought to be needed to accumulate high viral protein concentrations by driving efficient viral transcription and replication, so it is unclear how viruses can increase the concentration of viral proteins in early infection to allow IB formation. How viruses resolve this chicken-and-egg paradox to successfully establish infection is a major open question.

### Development of DARPin-P, a synthetic binder to RSV phosphoprotein

To visualize RSV infection establishment in real-time, we aimed to develop a live-cell imaging system with single vRNP sensitivity. We have recently shown for a different negative-strand RNA virus, influenza A virus, that single vRNPs can be visualized in living cells through expression of a fluorescently-labeled protein that binds to vRNPs in multiple copies^14^. As many copies of N and P proteins are bound to each RSV vRNP, we reasoned that, if a fluorescent protein could be developed that binds specifically to either N or P, such a protein would be recruited in many copies to a vRNP, yielding a sufficiently bright fluorescent signal to visualize single vRNPs during early infection^15,16^. Designed Ankyrin Repeat Proteins (DARPins) were assessed as potential N or P binders due to their small size (14-18 kDa), stability and low intracellular aggregation tendencies^17^. As a target, a previously described protein complex composed of the C-terminal domain of P (P_CTD_) associated with a NC-like structure consisting of rings of 10 to 11 N proteins (Nrings) was used^7,18^ (**Extended Data Fig. 1a**). High affinity DARPin binders were selected via ribosome display, from which one, DARPin-H6, was selected for further characterization based on its strong binding interaction (**Extended Data Fig. 1b**).

Mass spectrometry of DARPin-H6 pulldowns performed on RSV virion lysates, demonstrated that DARPin-H6 efficiently and selectively pulled down RSV P, with small amounts of other viral proteins also detected, suggesting DARPin-H6 binds directly to P in its native context (**Fig. 1a**). This interaction was further confirmed and localized to 40 residues at the N-terminal end of the P_CTD_ (aa 162-209) by 2D nuclear magnetic resonance (NMR) spectroscopy and gel shift assays using native agarose gel electrophoresis with recombinantly-expressed variants of N and P (**Fig. 1b, c and Extended Data Fig. 1c, d, e, f, g, h, i**). The binding kinetics of DARPin-H6 with P_CTD_ were assessed via biolayer interferometry (BLI), which showed a high affinity interaction with a dissociation constant (K_D_) of 12.3 nM (**Fig. 1d**). Based on the direct interaction with P, DARPin-H6 is referred to as DARPin-phosphoprotein (DARPin-P).

**Fig 1.**
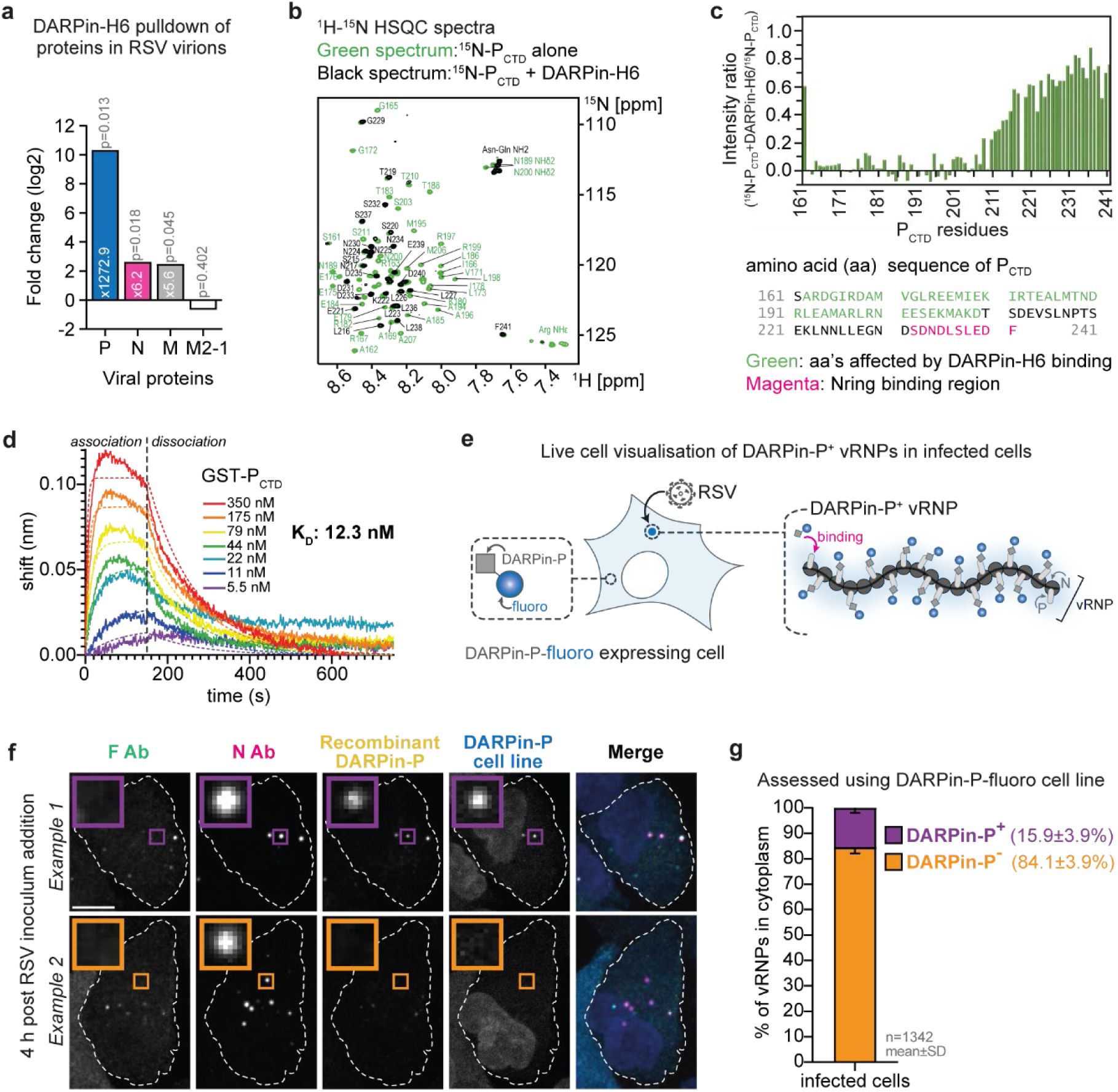
RSV vRNPs are heterogenous. **a,** Ribosome display identified DARPin-H6 as an RSV vRNP binder. DARPin-H6’s interaction with native RSV vRNPs were assayed by performing pulldowns on virion lysates followed by mass spectrometry. Fold change of detected viral proteins in the DARPin-H6 pulldown compared to beads only control is plotted. The adjusted P-value is recorded. **b, c,** NMR spectroscopy demonstrated that DARPin-H6 interacted with RSV P_CTD_. (**b**) ^1^H-^15^N HSQC spectrum of 50 µM ^15^N-P_CTD_ measured after addition of equimolar amounts of DARPin-H6 (black), superimposed on the spectrum of ^15^N-P_CTD_ alone (green). Assignments of signals that were fully broadened out by addition of DARPin-H6 are indicated in green font. (**c**) The intensities in the presence of DARPin-H6 divided by the intensities of the reference spectrum (intensity ratios) for each peak in the HSQC spectra are shown as a bar diagram. The amino acid (aa) sequence of P_CTD_ is shown below. Residues that are affected by DARPin-H6 binding are in green. Residues of the C-terminal Nring binding region are in magenta. **d**, Binding kinetics of the of P_CTD_ to DARPin-H6 was assessed using BLI with DARPin-H6 immobilized and P_CTD_ as analyte at indicated concentrations. Solid lines indicate original data and dashed lines indicate fitted curves. The DARPin-H6:RSV P_CTD_ interaction yielded a K_D_ value of 12.3 nM. Given its binding partner DARPin-H6 is hereon referred to as DARPin-phosphoprotein (DARPin-P). **e, f, g,** A549 cells expressing fluorescently-tagged DARPin-P (DARPin-P-fluoro), for live cell labeling of vRNPs with DARPin-P, were infected with WT RSV and stained for RSV F and N and with recombinant DARPin-P. Schematic (**e**), representative images (**f**) and quantification (**g**) are shown. Note that only cytoplasmic vRNPs (F^−^) were included in the quantification in (**g**). (**f**) Scale bars, 10 µm.

### DARPin-P labels a subset of vRNPs in infected cells

To assess the ability of DARPin-P to label individual vRNPs during infection, A549 cells were infected with wild-type (WT) RSV and stained with recombinant fluorescently-labeled DARPin-P together with fluorescently-conjugated anti-RSV F and N antibodies to mark vRNPs inside the host cell or in virions attached to host cells (**Extended Data Fig. 2a, b, c, d, e**). We identified single vRNPs that were strongly stained by DARPin-P, confirming that DARPin-P could be used for single vRNP detection (**Extended Data Fig. 2e**, *purple insert*). Surprisingly however, only a subset of vRNPs was labeled by the DARPin-P in infected cells (20±14%) (**Extended Data Fig. 2f**). Similarly, when a fluorescently-tagged DARPin-P (DARPin-P-fluoro) was expressed in cells (**Fig. 1e**), it robustly labeled only a subset of vRNPs in the cytoplasm of infected cells (16±4%) (**Fig. 1f, g and Extended Data Fig. 2g, h, i**), demonstrating the potential for live-cell single vRNP imaging by DARPin-P, and confirming that the heterogeneous labeling was not caused by post-fixation artefacts. Importantly, DARPin-P-fluoro expression in cells did not detectably interfere with RSV infection, as assessed by viral transcript and protein accumulation during infection (**Extended Data Fig. 2j, k, l, m**). Together, these results show DARPin-P-fluoro expressed in cells represents a powerful tool to visualize RSV infection with single vRNP resolution in living cells. Moreover, labeling of a subset of vRNPs by DARPin-P-fluoro suggests that vRNPs are heterogeneous and provides a technology with which to study vRNP heterogeneity.

### A subset of RSV virions contains DARPin-P^+^ vRNPs

To study the origin of vRNP heterogeneity and its consequence for infection, an additional vRNP labeling strategy is required that labels *all* vRNPs, which would allow comparison of infection by DARPin-P^−^ and DARPin-P^+^ vRNPs in live cells. Serendipitously, we found that cellular expression of exogenous, fluorescently-tagged P robustly labeled all vRNPs. To minimize the chance that the fluorescent tag on P would disrupt binding of exogenous P to vRNPs, we engineered exogenous P to encode a short peptide tag, SunTag^19,20^, and co-expressed a genetically-encoded antibody that binds to the SunTag peptide, called SunTag Antibody (STAb), fused to a fluorescent protein (referred to as P^exo^-fluoro) (**Fig. 2a**). P^exo^-fluoro labeled the large majority (93±4 %) of vRNPs in infected cells (**Fig. 2b, c and Extended Data Fig. 3a**), without detectably affecting viral transcription or replication (**Extended Data Fig. 3b, c, d, e, f**). Fortuitously, when the DARPin-P-fluoro and P^exo^-fluoro were co-expressed in cells and infected with RSV (**Fig. 2d**), DARPin-P still labelled only a small subset of vRNPs (19 ±11%), even though almost all vRNPs were associated with exogenous P and DARPin-P binds to P (**Fig. 3e**). These observations suggest that the DARPin-P signal is predominantly dependent on endogenous P, not P^exo^, possibly because P^exo^ levels on vRNPs are very low compared to endogenous P, or because the DARPin-P binds to a specific conformation of P present predominantly for endogenous P. Irrespectively, these results confirm that vRNPs exist in heterogenous states and that the DARPin-P-fluoro and P^exo^-fluoro tools combined allow real-time tracking and comparison of heterogeneous vRNPs in living cells.

**Fig 2.**
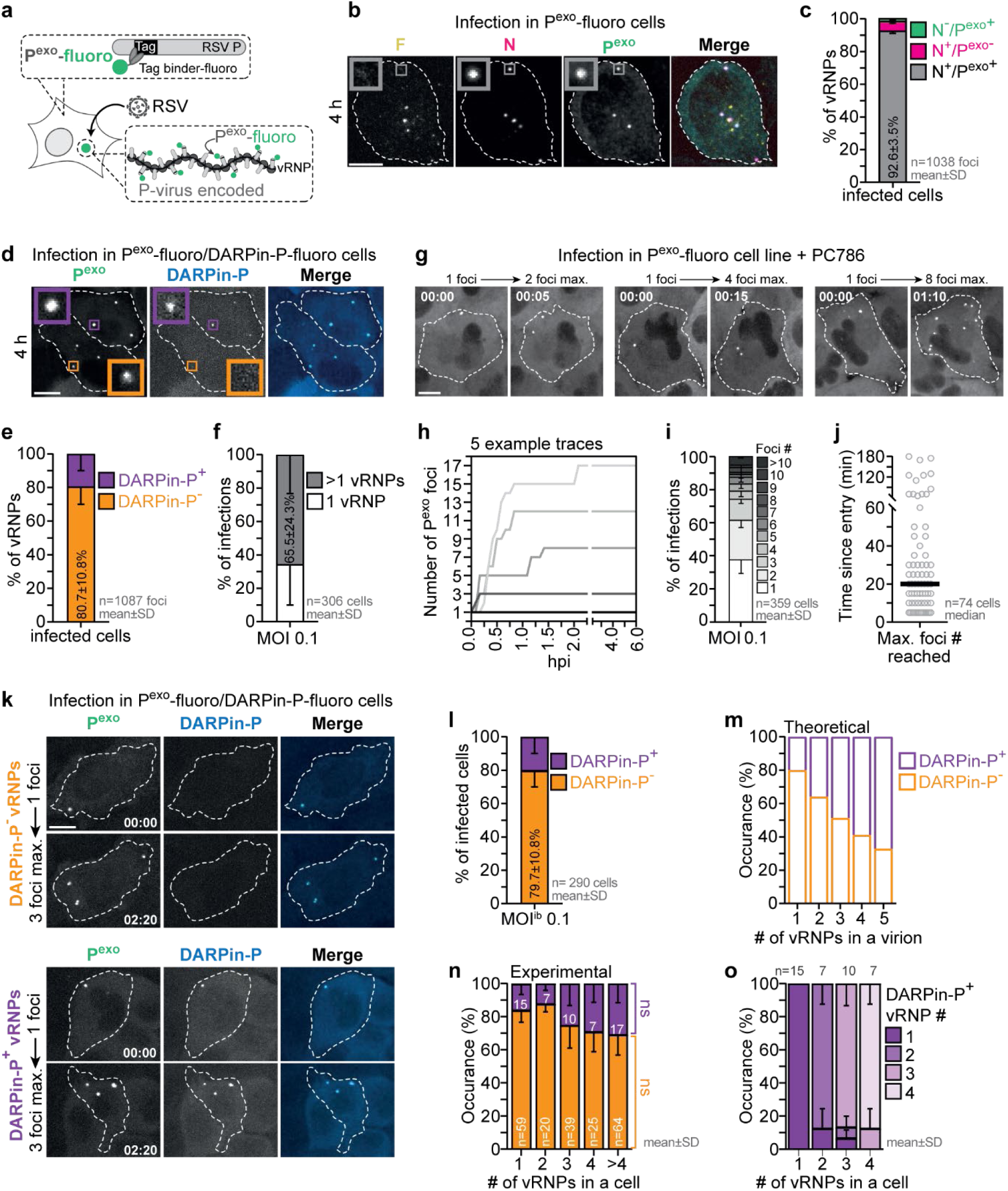
A subset of virions carry DARPin-P^+^ vRNPs. **a,** Schematic representation of the P^exo^-fluoro system for RSV vRNP visualization. **b, c,** P^exo^-fluoro foci colocalize with vRNPs (as labeled by anti-RSV N antibody). Anti-RSV F antibody staining was included to label intact virions, which are not labeled by P^exo^-fluoro. Representative images (**b**) and quantification of colocalization (**c**) are shown. **d, e,** Stable cell line expressing P^exo^-fluoro and DARPin-P-fluoro infected with WT RSV. Representative images (**d**) and quantification of colocalization (**e**) is shown. **f,** The number of vRNPs per infected cells at 4 hours post viral inoculum addition, is quantified. **g, h, i, j,** WT RSV infection in P^exo^-fluoro cells in the presence of the viral polymerase inhibitor, PC786. Representative images (**g**), example traces of the number of P^exo^-fluoro foci over time (**h**), quantification of the maximal number of vRNPs per cells (**i**) and the time taken for all virion containing vRNPs to separate (**j**) are shown. See also **Supplementary Video 1**. **k, l, m, n, o,** Infections by single virions were assessed to evaluate if all vRNPs originating from the same virion have the same DARPin-P labeling status. Representative images (**k**) and quantification (**l**) classifying infected cells with respect to their vRNPs DARPin-P status. If an infected cell had ≥1 vRNP that was DARPin-P^+^, the infected cells was classified as DARPin-P^+^. (**m**) Theoretical frequency of virions with at least one DARPin-P positive vRNP was calculated assuming DARPin-P^+^ vRNPs were randomly distributed over virions. (**n**) Experimental frequency of virions with at least one DARPin-P positive vRNP. Note that theoretical (**m**) and experimental data (**n**) do not match, indicating that DARPin-P^+^ vRNPs are not randomly distributed over virions. (**o**) Quantification of the fraction of vRNPs that are DARPin-P positive in each cell assigned as being infected with DARPin-P^+^ vRNPs in (**n**). Experiments were performed at low MOI (0.1) to ensure that the majority of cells were infected by a single virion. (**n**) Two-way ANOVA with Tukey’s multiple comparisons test compared if there was significant variation of the DARPin-P^+^ or DARPin-P^−^ infection fractions for each of the vRNP numbers. (**b, d, g, k**) Scale bar, 10 µm. (**g, k**) Time, h: min.

**Fig 3.**
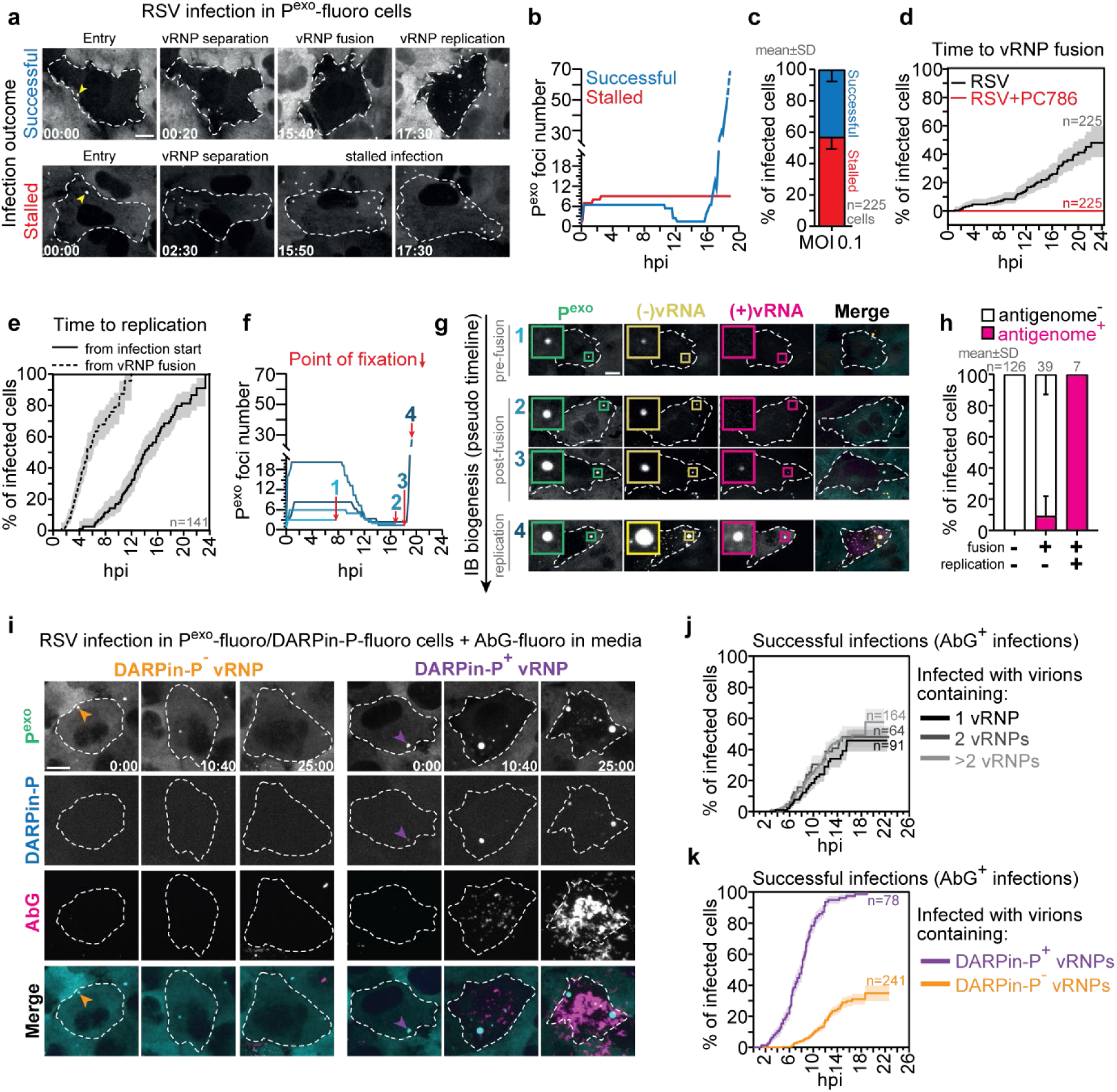
DARPin-P^+^ vRNPs give rise to successful infections. **a, b, c**, Time-lapse imaging of P^exo^-fluoro cells infected with WT RSV. Representative images from two time-lapse movies (**a**) and their P^exo^-fluoro foci count over time (**b**). Top image series show a successful infection and bottom image series show a stalled infection. Yellow arrowheads in (**a**) mark initial P^exo^-fluoro foci. See also **Supplementary Video 2**. (**c**) The frequency of successful and stalled infections is quantified. **d, e,** vRNP fusion and replication dynamics were evaluated in infection with >1 infecting vRNP. (**d**) Kaplan-Meier graph displaying the time to vRNP fusion relative to vRNP entry into the cell. No vRNP fusion is observed in the presence of PC786. (**e**) Time from vRNP entry (solid line) or from fusion (dashed line) to replication (as assessed by an increase in P^exo^-fluoro foci count). Note that vRNP fusion always precedes replication. **f, g, h,** Dynamics of vRNP fusion, IB formation and viral replication. Following P^exo^-fluoro time-lapse imaging, cells were fixed and genomes ((−)vRNA) and antigenomes ((+)vRNA) were stained by smFISH. (**f**) Time-traces of P^exo^-fluoro foci number for four representative infected cells, in which the point of fixation is noted by the red arrow. Representative images (**g**) and quantification (**h**) show the moment of antigenome appearance (a sign of replication) relative to vRNP fusion. **i, j, k,** Infection outcomes were assessed in relation to the infecting vRNP number and DARPin-P state. (**i**) Representative images of time-lapse movies of infections with a single DARPin-P^−^ (left) or DARPin-P^+^ (right) vRNP. Incoming DARPin-P^−^ and DARPin-P^+^ foci are indicated by orange and purple arrowheads, respectively. See also **Supplementary Video 3**. Kaplan-Meier graphs display infection success in relation to the number of infecting vRNPs (**j**) and to the infecting vRNPs DARPin-P status (**k**). Based on the vRNP activity status, hereon, DARPin-P^+^ vRNPs are referred to as Pre-Replication Centers (PRCs) and DARPin-P^−^ vRNPs as passive vRNPs. (**d, e, j, k**) Lines and shaded areas indicate mean and SE, respectively. (**a, g, i**) Scale bars, 10 µm. (**a, i**) Time, h: min.

Using the DARPin-P-fluoro and P^exo^-fluoro tools, we set out to characterize early RSV infection. Somewhat surprisingly, we found that many virions (66±24%) contained multiple vRNPs (**Fig. 3f**). To exclude that the multiple vRNPs observed in single cells at the onset of infection were a consequence of genome replication, infection were assessed in the presence of PC786, a potent RSV polymerase inhibitor that prevents vRNP transcription and replication^21^ (**Extended Data Fig. 3d, e, f**). For most infected cells, a single bright vRNP spot was observed upon viral entry, followed by a rapid increase in foci numbers (**Fig. 2g, h, Extended Data Fig. 3g and Supplementary Video 1**), demonstrating that multiple vRNPs enter together from a single virion and rapidly split after entry. Quantification of the number of vRNPs per infecting virion revealed that 38% contained 1 vRNP, 24% 2 vRNPs, 13% 3 vRNPs, 5% 4 vRNPs and 20% >4 vRNPs at an 0.1 MOI (**Fig. 2i**). While the majority of vRNPs originating from a single virion had split by 20 min post-entry, for virions containing large numbers of vRNPs this splitting process could take up to 3 h in rare cases (**Fig. 2j**). Therefore, we attribute any increase in vRNP number observed within the first 3 h of infection to the separation of vRNPs of the infecting virion. vRNA replication does not occur until >4 h after viral entry (see below, **Fig. 3e**), so vRNP splitting and replication can be reliably distinguished. While multiple vRNPs have been detected in RSV virions before by EM^22–24^, our results provide the first quantitation of vRNP copy number per virion, and reveal vRNP dissociation kinetics upon host cell entry.

Since many virions contained multiple vRNPs but only a subset of all vRNPs are DARPin-P positive, we asked whether individual virions typically contained either DARPin-P^+^ and DARPin-P^−^ vRNPs only, or mixed populations of vRNPs. Examining the fraction of DARPin-P^+^ vRNPs per virion revealed that DARPin-P^+^ vRNPs were not randomly distributed among virions, but rather that most virions contained either DARPin-P^+^ or DARPin-P^−^ vRNPs (**Fig 2k, l, m, n, o**). In summary, these results show that virions are heterogeneous in two different ways, they contain different numbers of vRNPs and their vRNPs can be in different states (DARPin-P^+^ or DARPin-P^−^).

### DARPin-P identifies pre-replication centers (PRCs) that drive successful infection

To assess functional consequences of virion heterogeneity on infection establishment and outcome, we followed single infected cells over time using the P^exo^-fluoro and DARPin-P-fluoro systems. First, the typical course of infection as observed by P^exo^-fluoro is described, after which the differences in the infection cycle between DARPin-P^+^ and DARPin-P^−^ vRNPs are detailed.

Infection with virions carrying >1vRNPs typically start with a single vRNP spot labeled by P^exo^-fluoro, which rapidly split into multiple foci (**Fig. 3a, b and Supplementary Video 2**), as discussed above (see **Fig. 2g, h, j**). In a subset of cells (43±8%) the intensity and size of one or more P^exo^-fluoro foci increased several hours after cell entry (**Fig. 3a, b, c, Extended Data Fig. 4a, b and Supplementary Video 2:** *successful infections*). Growing P^exo^-fluoro foci subsequently fused with other P^exo^-fluoro foci present in the same cell to form large, mostly immobile foci (**Fig. 3a**: *‘vRNP fusion’***, b, d, Extended Data Fig. 4a, b and Supplementary Video 2**). Typically, many new, smaller P^exo^-fluoro foci appeared after large P^exo^-fluoro foci were formed (**Fig. 3a**: *‘vRNP replication’***, b, e, Extended Data Fig. 4a, b and Supplementary Video 2**). Based on this series of events, we hypothesize that large P^exo^-fluoro foci formed by growth and vRNP fusion represent IBs, the sites of viral replication, and that the smaller P^exo^-fluoro foci formed later represent progeny vRNPs synthesized through viral replication. To test whether large P^exo^-fluoro foci indeed represent IBs, cells were fixed at the end of the time-lapse movie and large P^exo^-fluoro foci were assessed for viral RNA and proteins known to localize to IBs. Large P^exo^-fluoro foci stained strongly positive for viral genomes ((−)vRNA), antigenomes ((+)vRNA), viral transcripts and viral N and M2-1 proteins (**Fig. 3f, g and Extended Data Fig. 4c, d**), strongly suggesting that they represent IBs. Moreover, antigenomes, which are generated through viral replication, were exclusively observed in large P^exo^-fluoro foci formed after vRNP fusion, and not in smaller P^exo^-fluoro foci present before vRNP fusion (**Fig. 3f, g, h**), indicating that replication only occurs after large P^exo^-fluoro foci formation. Thus, we define large vRNPs that are formed through growth and fusion with other vRNPs as IBs, and define the moment of vRNP fusion as the moment of IB biogenesis. Further investigation into the new, small P^exo^-fluoro foci that appeared after IB formation, revealed that these foci stained positive for viral genomic RNA (**Extended Data Fig. 4e**), confirming that they represent progeny vRNPs. In all cases where vRNPs fused and IB’s were formed, vRNP progeny was also formed (**Fig. 3e**), indicating that viral replication occurs reliably upon IB formation. Moreover, since progeny vRNPs never formed before IB formation, we conclude that IB formation is prerequisite for vRNP replication and progeny production. While IB and vRNP progeny formation was observed in a subset of infections, in the majority of infections (57±8%), vRNP entry and initial splitting occurred normally, but IB formation failed to occur and no progeny vRNPs were produced (**Fig. 3a, b, c and Supplementary Video 2:** *stalled infections*). Taken together, these results show that IBs are nucleated by incoming vRNPs, that IB formation precedes viral replication and that IB nucleation represents the predominant bottleneck towards infection success.

To assess the effect of the number and DARPin-P status of incoming vRNPs on infection success, we combined the P^exo^-fluoro and DARPin-P-fluoro imaging systems (**Fig. 3i and Supplementary Video 3**). To simplify calling of ‘infection success’ (defined as infections that produce IBs and progeny vRNPs) in single infected cells, we made use of G protein staining at the plasma membrane, which we found is a simple and reliable proxy for IB and vRNP progeny formation (**Extended Data Fig. 4d, f, g**). G protein could be labeled and quantified in living cells using a non-neutralizing G antibody conjugated to a fluorophore (AbG-fluoro) added to the cell culture medium (**Extended Data Fig. 4h, i, j, k, l**). When examining the effect of vRNP number per virion on infection success, somewhat surprisingly, no difference was observed in infection success for virions with different numbers of vRNPs (**Fig. 3j**). We therefore turned our attention to DARPin-P status of incoming vRNPs. Virions containing DARPin-P^+^ vRNPs and DARPin-P^−^ vRNPs entered host cells with similar kinetics and showed similar vRNP numbers per virion (**Extended Data Fig. 4m, n**). However, in contrast to vRNP number, DARPin-P positivity was a very strong predictor of infection success: 100% of infections initiated by DARPin-P^+^ vRNPs were successful, while only 35% of infections originating from DARPin-P^−^ vRNPs were successful (**Fig. 3k and Extended Data Fig. 4o, p**). Furthermore, successful DARPin-P^+^ infections progressed faster than successful DARPin-P^−^ infections (**Fig. 3k and Extended Data Fig. 4o, p**). Interestingly, DARPin-P signal increased over time as DARPin-P^+^ vRNPs grew, with all IBs demonstrating especially strong DARPin-P staining (**Extended Data Fig. 4q**). Collectively, these results show that it is not the number of infecting vRNPs per virion that dictates the outcome of infection, but rather their DARPin-P state. Since DARPin-P stained both (a subset of) incoming vRNPs and IBs, and since DARPin-P^+^ incoming vRNPs went on to form IBs and successfully replicate to produce viral progeny, we refer to DARPin-P^+^ vRNPs as ‘pre-replication centers (PRCs)’. We refer to DARPin-P^−^ vRNPs as ‘passive vRNPs’, relating to their poor ability to drive infection success.

### PRCs undergo high rates of viral transcription

To understand why PRCs result in high rates of infection success, we examined if the transcriptional activity of PRCs and passive vRNPs were different during infection. To determine viral transcription rates in living cells, we made use of an assay to visualize single translating viral mRNAs, which we have previously developed^15^. In brief, an extra gene was introduced into the RSV genome that encodes for multiple repeats of the SunTag peptide epitope. Viral transcription produces mRNAs encoding for the SunTag array (SunTag mRNA), which are translated by host cell ribosomes. Upon translation, the SunTag peptides are co-translationally bound by the STAb-fluoro molecules expressed in host cells, resulting in bright fluorescent foci representing single translating mRNAs (**Fig. 4a**). The SunTag gene was inserted into the viral genome either between the viral P and M genes (upstream SunTag^eng^) or between the F and M2 genes (downstream SunTag^eng^) (**Extended Data Fig. 5a**). Introduction of the SunTag gene had minimal impact on viral growth kinetics, with comparable fitness to that of the previously described RSV-mCherry engineered strain^16^ (**Extended Data Fig. 5b, c**). Fluorescent SunTag mRNA foci appeared after infection with the SunTag^eng^ RSV strain. These represented translating SunTag mRNAs, as signals rapidly disappeared upon treatment with the translation inhibitor puromycin (**Extended Data Fig. 5d**). Importantly, the number of foci observed in live cells correlated very well with the number of SunTag mRNAs detected by smFISH in the same fixed cells (R^2^=0.97, **Extended Data Fig. 5e, f),** demonstrating that the number of translated SunTag mRNAs measured in live cells accurately reflects the number of total viral SunTag mRNAs, and can thus be used to determine viral transcription rates. Analysis of the number of SunTag mRNAs over time revealed that the rate of mRNA transcription was substantially higher in the SunTag^eng^ strain with the upstream insert compared to the SunTag^eng^ strain with the downstream insert, consistent with transcription rates being higher for genes at the 3’ end of the viral genome^25^ (**Extended Data Fig. 5g**). Together, these results show that the SunTag mRNA imaging method is a reliable and robust approach to quantify viral transcription dynamics in living cells.

**Fig 4.**
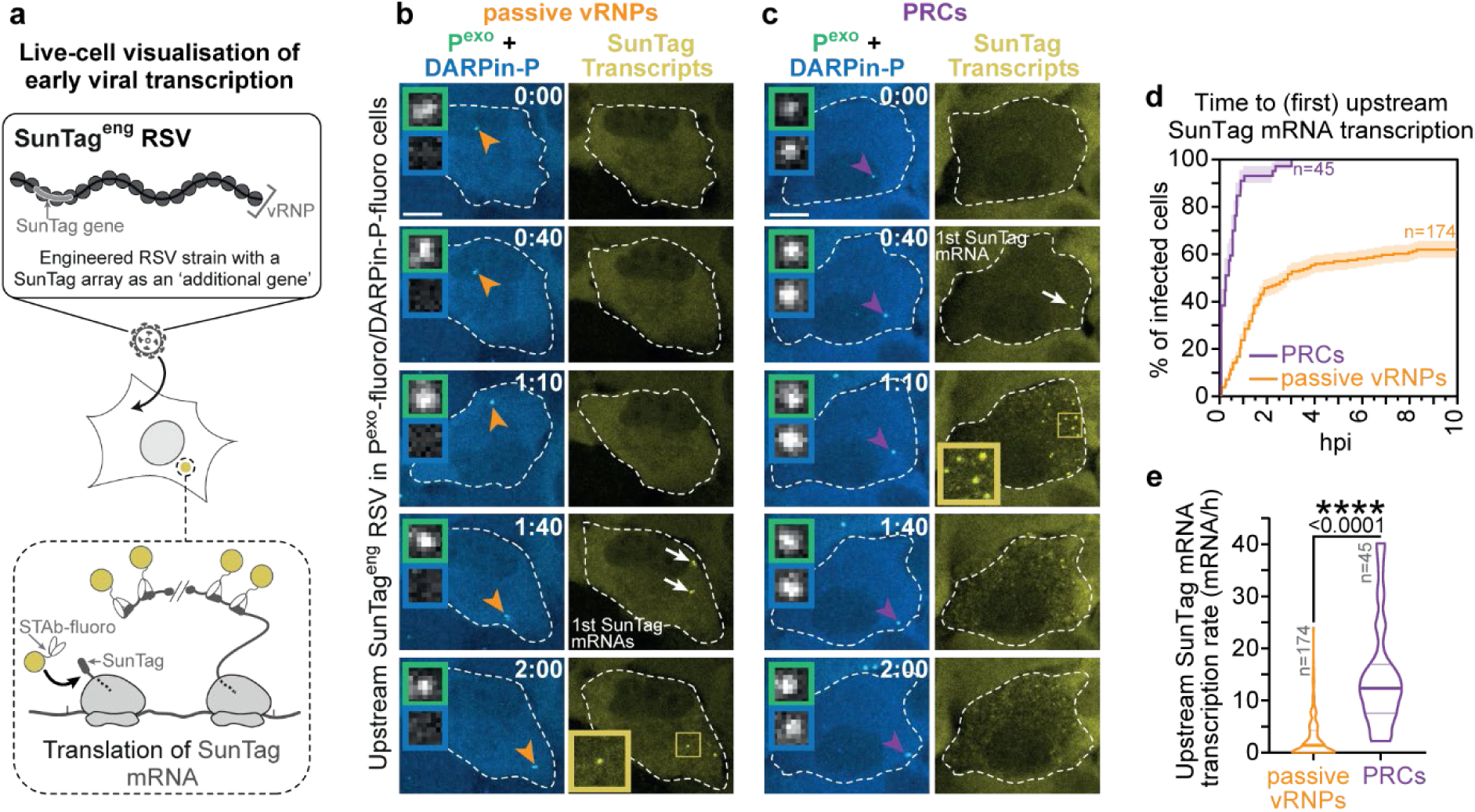
PRCs are highly active in transcription. **a, b, c, d, e,** An engineered RSV strain encoding a SunTag peptide array as an additional ‘viral gene’ (SunTag^eng^ RSV) provides a real-time readout of viral transcription. Here the SunTag array was inserted between the viral P and M genes (upstream SunTag^eng^ RSV). P^exo^-fluoro, DARPin-P-fluoro and SunTag mRNA imaging assays were combined in a single experiment. (**a**) Schematic of the SunTag imaging assay. Representative images of time-lapse movies of cells infected by either a passive vRNP (**b**) or PRC (**c**). Left image panel show a merge of P^exo^-fluoro and DARPin-P-fluoro signals while the right panel shows SunTag mRNA (STAb-fluoro foci) images. On the merged images, the orange and purple arrowheads identify passive vRNPs and PRCs, respectively, and the green and blue inserts show P^exo^ and DARPin-P signals, respectively, for the indicated vRNPs. The high translation rate of the PRC infection in (**c**) leads to the depletion of cellular STAb-fluoro due to cytoplasmic accumulation of mature SunTag peptides as observed from 1:40 onwards. See also **Supplementary Video 4**. (**d**) Kaplan-Meier graphs depict the start of viral transcription for infections by passive vRNPs and PRCs. Lines are mean, shaded areas are SE. (**e**) Viral transcription rate was calculated and plotted as violin plots with the median and quartiles shown (horizontal lines). Two-tailed unpaired Student’s t-test was used for statistical analysis. (**b, c**) Scale bars, 10 µm and time, h: min.

To interrogate viral transcription dynamics of heterogeneous vRNPs, we combined SunTag mRNA imaging with the P^exo^-fluoro and DARPin-P-fluoro imaging systems. The SunTag peptide used in the P^exo^-fluoro system to label P was replaced by an orthogonal tagging system, the ALFA-Tag and the nanobody-ALFA^26^, to enable combined P^exo^-fluoro and SunTag transcriptional imaging. Following three-color time-lapse imaging, transcriptional activity was assessed for both PRCs and passive vRNPs (**Fig. 4b, c, Extended Data Fig. 5h and Supplementary Video 4**). All infections with PRCs showed rapid viral transcriptional activation upon vRNP entry, with 100% of infection events showing transcription by 3 hpi, both for the upstream and downstream SunTag genes, compared to only 64% and 42% for passive vRNPs, respectively (**Fig. 4d and Extended Data Fig. 5i**). Even for the passive vRNP infections that did show transcription, transcriptional dynamics were severely delayed, with only 28% of the upstream and 11% of the downstream infections showing transcripts at 1 hpi while 94% of upstream and 50% of downstream PRC infections showed transcripts at that time-point (**Fig. 4d and Extended Data Fig. 5i**). PRC-containing infections also had substantially higher rates of transcription (5.4x and 15.1x higher for the upstream and downstream SunTag^eng^ RSV strains, respectively) than their passive vRNP-containing counterparts (**Fig. 4e and Extended Data Fig. 5j**). Collectively, these findings demonstrate that PRCs have much higher rates of transcription, likely explaining their high infection success.

### PRCs contain high levels of viral polymerase-complex proteins

Since PRCs have increased transcriptional activity compared to passive vRNPs, it is possible that their molecular composition differs. To test this, we undertook a molecular analysis of vRNP composition and asked whether vRNP compositional heterogeneity could explain differences in infection progression. Interestingly, in time-lapse movies of single vRNPs, we noticed that PRCs appeared to move somewhat more slowly than passive vRNPs, suggesting that they were of larger size. Indeed, rapid kinetic measurements revealed that PRCs show a ∼2x lower diffusion rate than passive vRNPs (0.032 vs 0.063 µm^2^/sec for PRCs and passive vRNPs, respectively) (**Extended Data Fig. 6a, b**), but not as slow as the diffusion rate of IBs (0.004 µm^2^/sec) (**Extended Data Fig. 6a, b**). These results suggest that PRCs may represent an intermediate in between vRNPs and IB-like structures, potentially explaining their high transcription and replication activity.

The slower diffusion suggests that PRCs may be associated with higher viral protein and/or RNA copy numbers. We first assessed viral genome and antigenome content of PRCs and passive vRNPs by smFISH, which showed that both subsets of vRNPs contained mostly genomes, with a low frequency of vRNP foci containing both a genome and an antigenome (**Fig. 5a and Extended Data Fig. 6c**). Interestingly, the genome smFISH staining for PRCs was 1.6-fold higher in intensity than their passive vRNP counterparts (**Fig. 5b**), suggesting that a subset of PRC foci contains two or more genomes (discussed in more detail later). We conclude that PRCs have a slightly higher genome content than passive vRNPs, but that this difference in genome content is substantially smaller than the difference in transcriptional activity (>10-fold), demonstrating that individual PRC genomes have much higher transcriptional activity than genomes of passive vRNPs.

**Fig 5.**
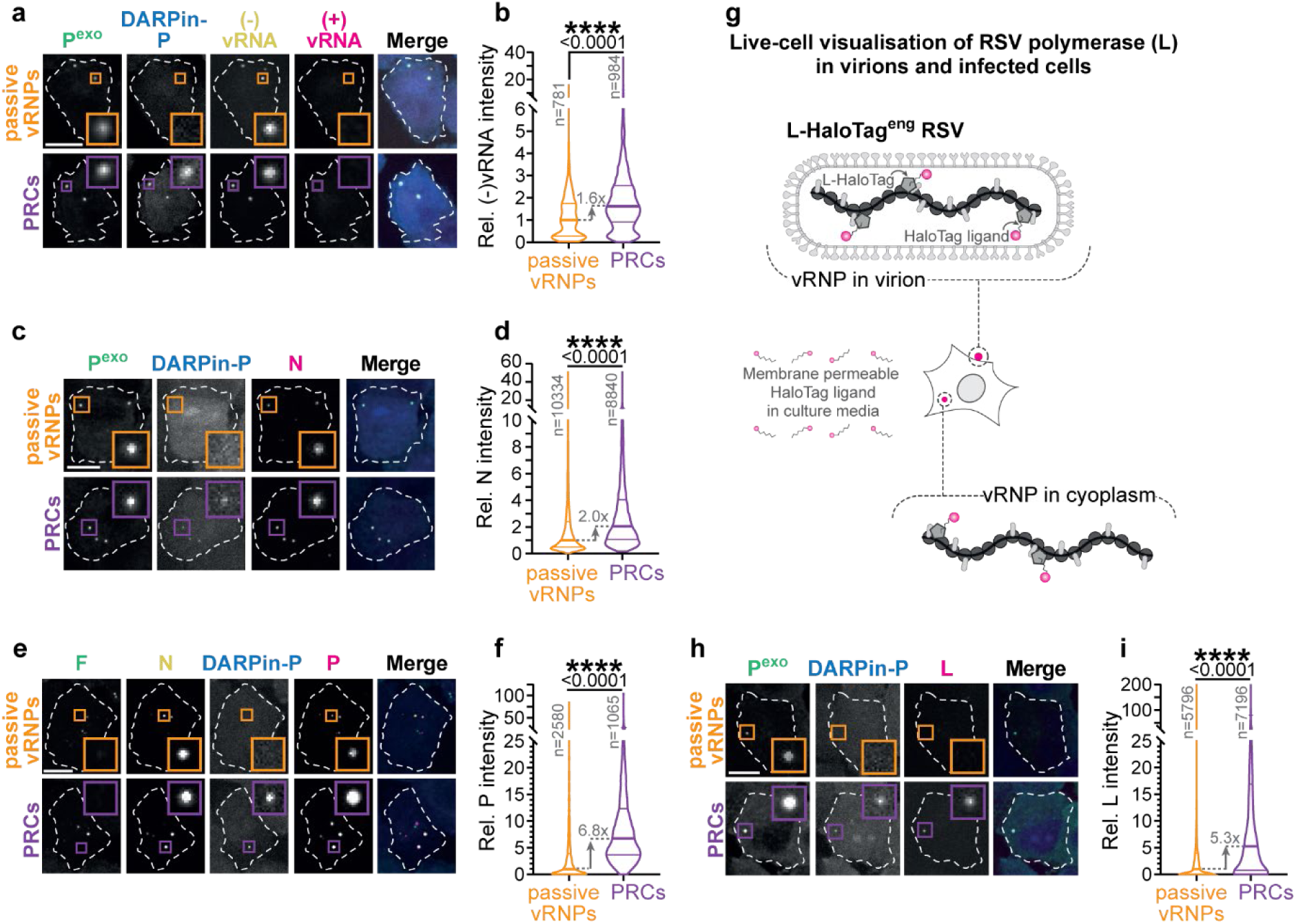
PRCs have higher quantities of associated viral proteins. Incoming passive vRNPs and PRCs were assessed at 4 hours following viral inoculum addition. The translation inhibitor emetine was added to cells together with virus addition to prevent progression of infection. **a, b,** genome ((−)vRNA) and antigenome ((+)vRNA) presence in vRNPs was evaluated. (**a**) Representative images of smFISH staining carried out on the P^exo^-fluoro/DARPin-P-fluoro cell line is shown. (**b**) Quantification of the (−)vRNA intensity is shown for passive vRNPs and PRCs. **c, d, e, f, g, h, i,** RSV vRNP-constituent protein levels were assessed in passive vRNPs and PRCs. Representative images (**c**) and quantification (**d**) of nucleoprotein (N) levels. Representative images (**e**) and quantification (**f**) of phosphoprotein (P) levels. (**g**) Schematic highlighting the L-HaloTag^eng^ RSV strain which enables real-time visualization of viral polymerase levels in the presence of fluorescently-tagged, cell membrane-permeable HaloTag ligands. This strain allows visualization of viral polymerase levels both in the virion and on vRNPs in infected cells. Representative images (**h**) and quantification (**i**) of L levels. (**b, d, f, i**) Violin plots show median and quartiles (horizontal lines). (**b, d, f, i**) Two-tailed unpaired Student’s t-test was used for statistical analysis. (**a, c, e, h**) Scale bar, 10 µm.

To further explore compositional variations in PRCs and passive vRNPs that could result in their functional differences, we assessed the levels of viral N, P and L, the essential viral polymerase-complex proteins. vRNAs are thought to be fully encapsidated by N proteins^6,7,27^, and as such we would expect N levels to mirror the vRNP content. Using antibody staining we found that PRCs were associated with 2-fold higher N levels than passive vRNPs (**Fig. 5c, d**). The slightly higher levels of N protein compared to the genomic content (2.0-fold vs 1.6-fold) was highly reproducible and may reflect additional free N, referred to as N^0^, associated with PRCs. N^0^ is known to associate strongly with IBs, so association of small amounts of N^0^ with PRCs is consistent with these vRNPs representing IB precursors.

We next examined P levels on single vRNPs. Since DARPin-P binds to RSV P protein, higher levels of P on PRCs were expected. Indeed, PRCs contained 6.8-fold higher P levels than passive vRNPs (**Fig. 5e, f and Extended Data Fig. 6d**). We next assessed L protein (viral polymerase) levels on incoming RSV vRNPs. As there are no commercially available L antibodies, we generated an engineered RSV strain carrying a single HaloTag in the coding sequence of the L gene, which can be labelled by membrane-permeable fluorescent dyes^28^ (L-HaloTag^eng^ RSV, **Fig. 5g and Extended Data Fig. 6e**). L levels were 5.3-fold higher in intensity on PRCs than passive vRNPs (**Fig. 5h, i and Extended Data Fig. 6f**). The levels of P and L proteins associated with PRCs (5-7-fold) are substantially higher than the difference in the number of genomes (1.6-fold), indicating that PRCs have a higher viral protein occupancy *per genome*. The higher levels of viral polymerase-complex proteins on PRCs provide a plausible explanation for their high transcriptional activity and more rapid replication dynamics.

### PRCs represent pre-assembled biomolecular condensate seeds that nucleate IBs

We considered two, non-mutually exclusive explanations for the substantially higher levels of viral polymerase-complex proteins associated with PRCs; first, it is possible that PRCs originate from virions with higher viral protein levels. Second, PRCs may show stronger binding to viral proteins, retaining them better upon cytoplasmic entry. To assess these two possibilities, we measured L levels both in virions and subsequently on single vRNPs after cytoplasmic entry using the L-HaloTag^eng^ strain (**Fig. 5g and 6a**). We found that L intensity was not just higher on cytoplasmic PRCs, but also in the virions from which the PRCs originated from (**Fig. 6b**), demonstrating that vRNP compositional heterogeneity originates, at least in part, from infecting virions.

**Fig 6.**
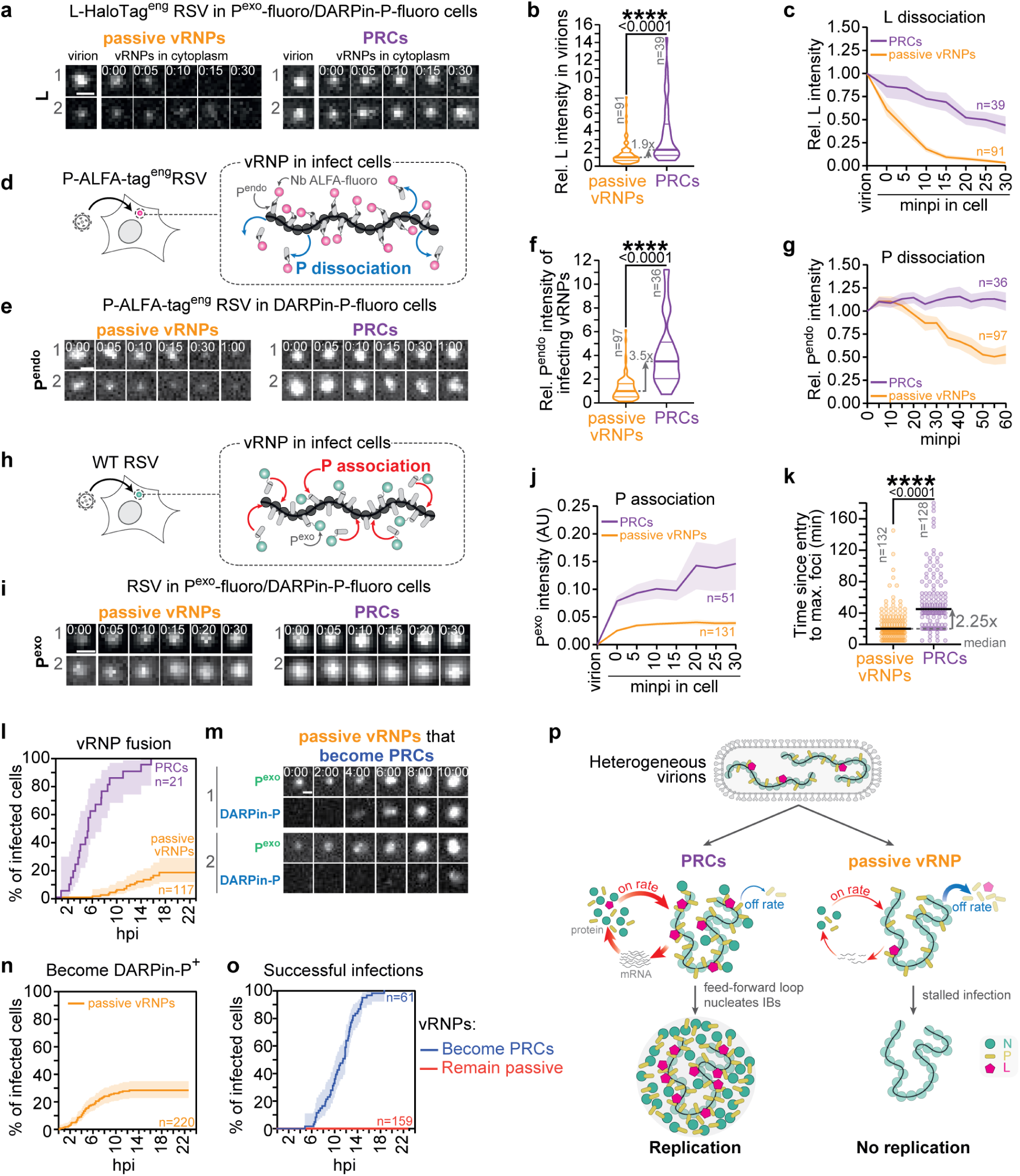
PRCs are biomolecular condensate seeds. **a, b, c,** Visualization of L protein levels in virions and subsequently on vRNPs after host cell viral entry. L protein is visualized using the L-HaloTag^eng^ RSV strain. Infections are carried out in the P^exo^-fluoro/DARPin-P-fluoro cell line. (**a**) Two representative image series highlighting the L protein levels in the virion prior to viral entry and vRNP following viral entry for passive vRNPs and PRCs. Quantification of L levels in virions (**b**) and L levels on vRNPs over time (**c**) are shown for virions containing passive vRNPs and PRCs. **d, e, f, g,** Endogenous P (P^endo^) associated with vRNPs was visualized using the P-ALFA-Tag^eng^ RSV strain. (**d**) Schematic highlighting how the P-ALFA-Tag^eng^ RSV strain allows visualization of endogenous viral P protein upon infection and vRNP release into the cytoplasm of cells expressing Nb-ALFA-fluoro. Using cells that additionally express DARPin-P-fluoro allows establishment of vRNP’s activity status. (**e**) Two representative image series of vRNP P^endo^ signal over time in passive vRNPs and PRCs. Quantification of P^endo^ levels on the infecting vRNP (**f**) and the relative P^endo^ levels on vRNPs over a 1 h time frame after viral entry (**g**) for passive vRNPs and PRCs is shown. **h, i, j,** Viral protein association kinetics of passive vRNPs and PRCs were assessed using the P^exo^-fluoro/DARPin-P-fluoro system. Infections were carried out using the P-ALFA-Tag^eng^ RSV strain in cells that additionally expressed Nb-ALFA-fluoro, and used to identify infecting vRNPs independent of the P^exo^-fluoro signal. (**h**) Schematic highlighting how infections in P^exo^-fluoro cells allow visualization of exogenous viral P protein association on cytoplasmic vRNPs. (**i**) Two representative image series of vRNP P^exo^-fluoro signal over time in passive vRNPs and PRCs. (**j**) Quantification of the P^exo^-fluoro levels on vRNPs over a 30 min time frame. **k,** Infecting vRNP separation kinetics were quantified with respect to the vRNPs activity state. **l,** Kinetics of vRNP fusion is displayed as Kaplan-Meier graphs, separated in accordance to the vRNP’s activity state. **m, n, o,** Infections with incoming passive vRNPs were further assessed. (**m**) Two representative image series from time lapse movies highlight passive infecting vRNPs that become PRCs during the course of infection. See also **Supplementary Video 5**. (**n**) Kaplan-Meier graphs showing the kinetics of passive vRNPs becoming PRCs during infection. (**o**) Kaplan-Meier graphs showing the success of passive vRNPs that either do or do not become PRCs during infection. **p,** Schematic highlighting the mechanism by which a subset of infecting vRNPs go onto seed IBs. (**c, g, j, l, n, o**) Lines are mean and shaded region SE. (**b, f, k**) Two-tailed unpaired Student’s t-test was used for statistical analysis. (**a, e, i, m**) Scale bars, 1 µm and time, h: min.

When L intensity on single vRNPs was followed as virions fused with host cells and released their vRNPs into the cytoplasm, a substantial drop in L intensity was found when comparing virions with their vRNPs post-entry, suggesting that a substantial fraction of L protein dissociate from vRNPs post-entry (**Fig. 6a, c**). Interestingly, while the level of L protein associated with the vRNPs immediately post-entry was 1.4-fold higher for PRCs than for passive vRNPs, the difference in L levels dramatically increased to 13-fold over a 30 min time frame (**Fig. 6c**), demonstrating that L rapidly dissociated from passive vRNPs, while remaining tightly bound to PRCs. To determine if P shows a similar higher off-rate from passive vRNPs compared to PRCs after cell entry, we engineered an RSV strain in which the endogenous P protein was fluorescently-tagged (referred to as P^endo^, see Methods) (**Fig. 6d**). The P^endo^ strain demonstrated growth kinetics comparable to WT RSV (**Extended Data Fig. 7a**), and allowed robust detection of endogenous P levels, as 98±1.% of virions contained detectable levels of the tagged P protein (**Extended Data Fig. 7b, c**). The levels of endogenous P on incoming PRCs vRNPs was 3.5-fold higher than on passive vRNPs, consistent with our antibody staining (**Fig. 6e, f and see 5e, f**). Interestingly, P protein rapidly dissociated from passive vRNPs, as was observed for L, while remaining tightly associated with PRCs, suggesting that the P affinity for vRNPs, like is the case for L, depends on their vRNP status (**Fig. 6e, g**). We next explored association kinetics of P towards PRCs and passive vRNPs. We examined binding kinetics of P^exo^-fluoro to vRNPs immediately after viral entry into the cytoplasm (**Fig. 6h**). We observed that PRCs not only brought in more endogenous P (assessed via P^endo^), but also attracted more P that was present in the cytoplasm (assessed via P^exo^-fluoro binding) compared to passive vRNPs (**Fig. 6i, j**). The substantially higher levels of both endogenous P and P^exo^-fluoro on PRCs compared to passive vRNPs likely explains why PRCs are selectively labeled (above the detection threshold) by DARPin-P-fluoro (which binds to P), even though all vRNPs are bound by P^exo^-fluoro. In addition to P levels, it is possible that DARPin-P also preferentially binds to PRCs because of conformational differences of P bound to passive vRNPs or PRCs. Our NMR data shows that DARPin-P binds to a structured conformation of the P_CTD_ (see **Fig. 1b, c and Extended Data Fig. 1e, f**), and we cannot exclude that this conformation of P is more abundant on PRCs. It is noteworthy that a structured conformation of P_CTD_ is known to bind L^29^, and may thus represent an ‘active’ conformation. It is therefore possible that PRCs have more active P, and that this further contributes to DARPin-P selectively towards PRCs. In conclusion, PRCs have lower off-rates and increased attraction of viral proteins (on-rates), indicating that they are more efficient at retaining and recruiting viral proteins (i.e. they are ‘sticky’ for viral proteins), likely explaining their potent ability to nucleate biomolecular condensates (i.e. IBs) which are formed through multimeric viral protein-protein interactions.

IBs not only display high concentrations of viral proteins, but also of viral RNAs^1^. We therefore assessed whether viral genomic RNAs (i.e. vRNPs) also show increased binding affinity for PRCs. First, we measured the separation dynamics of vRNPs originating from a single virion upon cytoplasmic entry, and observed that on average PRCs took significantly longer to split compared to passive vRNPs (45 min vs 20 min) (**Fig. 6k**). This result indicates that PRCs are not only more effective at recruiting and retaining viral proteins, but also show increased affinity towards other vRNPs. Possibly, the affinity to other vRNPs is so high, that a subset of incoming vRNPs remain together without splitting, providing a plausible explanation for why PRCs have a slightly higher average viral genomic RNA content than passive vRNPs (See **Fig. 5b**). The increased affinity of PRCs for other vRNPs was also observed during later infection, where 100% of PRCs underwent vRNP fusion, with a median time of 5.5 hpi (**Fig. 6l**). In contrast, only 16% of passive vRNPs showed vRNP fusion, and the fusion events that did occur, happened substantially later in infection (**Fig. 6l**). These observations show that PRCs recruit not only viral proteins, but also other vRNPs. It is likely that most incoming PRCs are not yet ‘sticky’ enough to maintain vRNP association, explaining initial vRNP splitting, but as PRCs grow due to new viral protein synthesis and association, their ‘stickiness’ increases with the increasing size of the protein-protein interaction network to the point that vRNPs start to fuse (**Extended Data Fig. 7d**). Since many of the features that define mature IBs are already present in incoming PRCs, PRCs resemble biomolecular condensate ‘seeds’ that are primed to nucleate IBs.

If PRCs indeed seed IBs we would predict that PRCs would always precede IB formation. Our earlier results showed that a subset of incoming passive vRNPs eventually undergo viral replication (see **Fig. 3k**: *DARPin-P^−^ vRNPs*), and we wondered whether vRNPs in these infections transitioned from passive vRNPs to PRCs (i.e. became PRCs) before undergoing IB formation and subsequent replication. Data demonstrating that all mature IBs were DARPin-P^+^ (see **Extended Data Fig. 4q**) provided preliminary evidence that a subset of passive vRNPs could transition into PRCs, and eventually develop into mature IBs. Indeed, we observed that in all cases where incoming vRNPs started as passive vRNPs but did successfully form IBs (30% of infections starting with passive vRNPs), one or more passivevRNP eventually became PRCs at a later stage during the infection process, but always before viral replication ensued (**Fig. 6m, n and Extended data Fig. 7e, f**). It was specifically the infections where one or more vRNP became PRCs that subsequently went on to successfully establish IBs (**Fig. 6o**). No infections were successful without vRNPs first acquiring the PRC-status. When we examined infections for which passive vRNPs eventually acquired the PRC state, they showed higher transcription rates even during very early infection compared to passive vRNPs that remained passive (**Extended data Fig. 7g, h, i, j**). Moreover, these passive vRNPs that went on to become PRCs, also demonstrated increased recruitment and retention for viral proteins (**Extended data Fig. 7k**). These results suggest that DARPin-P positivity may not be a binary phenotype, but rather a continuum that reports on the degree to which incoming vRNPs are in a PRC-state.

## Discussion

In this study we have developed technologies to visualize RSV infection with single vRNP resolution in living cells. Using these technologies, we have tracked viral infection from vRNP entry through viral transcription, IB formation and viral replication. We identify PRCs as crucial structures that nucleate IBs and are thus essential for viral genome replication. Our results identify two key features of PRCs that make them potent nucleators of IBs. First, PRCs are transcriptionally highly active, driving rapid accumulation of newly-synthesized viral proteins needed for vRNP growth into IBs. Second, PRCs have high affinities for viral proteins, resulting in high stability at low viral protein concentrations and efficient recruitment of newly synthesized viral proteins, features essential to nucleate biomolecular condensates at low viral protein concentrations. Together, these features result in a feed-forward loop that drives rapid growth of PRCs into IBs (**Fig. 6p**). Importantly, we find that PRCs are pre-formed and packaged into virions, which allows rapid kickstarting of viral infection once vRNPs enter host cells. Since only a small subset of virions contain PRCs (∼20%), our results also identify virion heterogeneity as a major cause of cell-to-cell variation in RSV infection.

### RSV virions are compositionally diverse

We observed virion diversity with respect to the number and type of vRNPs present. While previous work has shown that multiple vRNPs can be packaged into single virions^22–24^ our work has provided a quantitative description of this phenomenon. More importantly, our study has uncovered the functional consequences of carrying multiple vRNPs in individual virions. Surprisingly, we find that the number of vRNPs per virion is not a major determinant of infection success, but rather it is the vRNP-state which determines infection success. Interestingly, vRNPs originating from the same virion are generally in the same vRNP-state. Therefore, a subset of RSV virions, those containing one or more PRCs, are highly infectious and most likely the main drivers establishing a successful (primary) infection. Why certain virions contain mostly PRCs while others contain mostly passive vRNPs, remains an interesting open question. One possibility is that virion composition is determined at the level of virus-producing cells. For example, PRCs may be produced by late-stage infected cells when viral proteins are abundant in virus-producing cells, whereas passive vRNP containing virions are produced early in infection when viral proteins are present at lower levels. Alternatively, the subcellular localization of virion budding may determine the type and number of vRNPs that are packaged.

### Temporal coordination of viral transcription and replication

The RSV genome acts as a template for both viral transcription and replication. Viral transcription should precede viral replication, because newly synthesized N protein is required for encapsidation of new genomes and antigenomes produced during replication. However, if and how transcription and replication are temporally coordinated remained unknown. Here, we show that even though transcription occurs in IBs during late-stage infection, IBs are not *required* for viral transcription, as viral transcription occurs on vRNPs that are freely diffusing in the cytoplasm, before the formation of IBs. In contrast, our results show that replication occurs exclusively in IBs, and as such, IBs are essential for viral replication. Since IB formation requires synthesis of new viral proteins and thus requires viral transcription, IB formation temporally coordinates viral transcription and replication to ensure correct vRNA encapsidation.

### Biogenesis of viral biomolecular condensates

IBs have been demonstrated to concentrate the RSV vRNP constituent proteins N, P, L and M2-1^1,11,30–32^, as well as viral genomes and transcripts^1,33–35^, and constitute key sites of viral transcription and replication in late stage viral infection. IB formation is likely driven by low-affinity multimeric protein-protein interactions mediated by the intrinsically disordered regions (IDRs) of N and P^36–39^. Despite our understanding of the composition and physiochemical properties of IBs, the mechanism of their biogenesis during natural infection was lacking. Here, we show that IBs are typically nucleated by individual vRNPs, rather than being formed *de novo* in the cytoplasm. Using vRNPs as nucleation seeds likely reduces the critical concentration of cytoplasmic viral protein needed to drive condensate nucleation, and as such may enable IB formation during early infection when viral protein concentrations are low. PRCs are potent nucleators of IBs, since they are already associated with high levels of viral N, P and L protein. High levels of viral polymerase complex proteins likely result in higher viral transcription and protein synthesis rates, which results in recruitment of even more viral proteins to these vRNPs. More importantly, higher levels of N and P in PRCs likely also create a stable multimeric protein interaction network surrounding vRNPs that prevents N, P and L molecules from dissociating from PRCs upon host cell entry, and creates increased numbers of binding sites to recruit newly-synthesized viral N, P and L proteins, further enhancing their growth into IBs. Higher transcription rates combined with higher rates of viral protein recruitment create an effective positive feedback loop that drives PRC growth into IBs. Once viral protein interaction networks in PRCs have grown above a critical threshold, PRCs can either fuse with other vRNPs, if present in infected cells, or themselves form mature IBs. The feedback loop that drives PRC growth can also result in stalled infections when vRNPs associated with low levels of viral proteins enter the host cell. Not only do these vRNPs produce few viral proteins due to low transcription rates, but viral proteins also dissociate rapidly from these vRNPs. All viruses of the order *MNV* encode N, P and L proteins carrying out conserved functions^40^, it is therefore tempting to speculate that the IB nucleation and growth mechanisms discovered here may reflect a common mechanism for viruses of the *MNV* order. More broadly, our study on IB nucleation may provide a paradigm for condensate nucleation in general.

## Methods

### Cell lines and cell culture

A549 (ATCC, cat. no. CCL-185), BSR T7/5^41^ (BHK-21 cells that constitutively express the T7 RNA polymerase) and HEK293T cells (ATCC, cat. no. CRL-3216) were cultured in DMEM (Thermo Fisher Scientific, cat. no. 31966021) supplemented with 10% fetal bovine serum (FBS) (Merck, cat. no. F7524) and 1% penicillin-Streptomycin (Pen-Strep) (Thermo Fisher Scientific, cat. no. 15140122). Vero cells (ATCC, cat. no. CCL-81) were cultured in DMEM supplemented with 5% FBS and 1% Pen-Strep. HEp-2 cells (ATCC, cat. no. CCL-23) were cultured in MEM (Thermo Fisher Scientific, cat. no. 42360032) with 10% FBS and 1% Pen-Strep. Cells were cultured at 37°C and 5% CO_2_. Cell lines used in this study were confirmed to be mycoplasma negative.

### RSV protein samples

RSV Nrings without tag were purified by co-expression in *E. coli* and co-purification with GST-P_CTD_ (aa 161-241), using the GST-tag, as described previously^18^. When specified, the GST tag was removed by thrombin cleavage. For NMR measurements, ^15^N-labeled P_CTD_, cloned into the pGEX-4T3 plasmid, were used to express and purify ^15^N-labeled P_CTD_ constructs from *E. coli* as previously described^42^. For expression and purification of all the GST-P fragments, the corresponding P sequences were cloned into pGEX-4T3 plasmid and the proteins were expressed in *E.coli*. Purifications were performed as previously described^18^.

### Selection and screening of DARPins binding to RSV vRNP

To generate DARPin binders for the GST-P_CTD_+Nrings complex, biotinylated GST-P_CTD_+Nrings complex was immobilized alternatingly on either MyOne T1 streptavidin-coated beads (Thermo Fisher Scientific, cat. no. 65601) or Sera-Mag neutravidin-coated beads (Cytiva, cat. no. 78152104011150) depending on the selection round. Ribosome display selection was performed as previously described^43^, but using a semiautomatic KingFisher Flex MTP96 well platform.

The fully synthetic library includes N3C-DARPins with three randomized internal repeats with the original randomization strategy as reported^44^, but including a stabilized C-cap^17,45,46^. Additionally, the library is a 1:1 mixture of DARPins with randomized and non-randomized N- and C-caps, respectively^17,47^, and successively enriched pools were ligated in a ribosome display-specific vector^43^. Selection was performed over four rounds with decreasing concentrations of biotinylated GST-P_CTD_+Nrings complex (250 pmol, 125 pmol, 5 pmol) for the first three rounds and 50 pmol of target for the last recovery round, and increasing washing steps^43,48^. For rounds 2 to 4, prepanning with biotinylated GST was used to remove potential binders against GST.

To screen individual DARPins for their binding properties, the selected pool of DARPins from ribosome display was subcloned by restriction digest with BamHI and HindIII into the pQE30-derived bacterial expression vector pQIq (Qiagen). This creates DARPins with an N-terminal MRGS(H_6_)-tag and a C-terminal FLAG-M2-tag. 192 single DARPin clones were screened against GST-P_CTD_+Nrings complex and GST only, both directly immobilized, by a crude extract ELISA. The crude extracts were prepared as described previously^49^. 32 identified DARPin clones were sequenced and 13 of them were unique in their sequence. These 13 clones were IMAC purified and validated in an ELISA, described previously^49^, against GST-P_CTD_+Nrings complex and GST.

### Purified DARPin-P production

DARPin-P-H6 with an N-terminal MRGS(H_6_)-tag and a C-terminal FLAG-M2-tag (His_6_-DARPin-P-H6-FLAG, referred to as DARPin-H6 in **Fig. 1** and DARPin-P thereafter, originally obtained as 011-1055-C6-2605-H5 in the selection, was cloned into the pQIq vector backbone^50^ (**Supplementary Table 1**). The plasmid was further modified to generate fluorescent-protein fusions, DARPin-P-BFP and DARPin-P-mRuby3 (**Supplementary Table 1**). DARPin-P and the fluorescent fusion proteins were recombinantly expressed in *E. coli* XL-1 Blue cells (Agilent, cat. no. 200249) and purified using the MRGS(H_6_)-tag. In brief, a 20 ml primary culture of transformed cells was grown overnight in 2x YT medium (Merck, cat. no. Y2377-250G) supplemented with 1% glucose and 100 µg/ml ampicillin (Merck, cat. no. A5354-10ML) at 37 °C with shaking at 160 rpm. The primary culture was diluted into 400 ml of the same medium and incubated at 37 °C with shaking at 160 rpm until the OD_600_ reached 0.5–0.8. Protein expression was induced with 0.5 mM IPTG (Thermo Fisher Scientific, cat. no. AM9464), and the culture was further incubated at 37 °C and 160 rpm for 4 h. Cells were harvested by centrifugation at 4000 × g for 20 minutes and stored at −20 °C until use. The cell pellets were thawed and resuspended in 25 ml of buffer A (PBS, pH 7.2, supplemented with 150 mM NaCl, 30 mM imidazole) containing 5% [v/v] glycerol and cOmplete Mini EDTA-free Protease Inhibitor Cocktail (Merck, cat. no. 11836170001). Cells were lysed by sonication, and the lysate was clarified by centrifugation at 20,000 × g for 30 minutes. The resulting clear supernatant was loaded onto an Ni-NTA agarose column (Thermo Fisher Scientific, cat. no. R90115), which was subsequently washed with 20 column volumes (CV) of buffer A. Bound MRGS(H_6_)-tagged proteins were eluted using buffer A supplemented with 400 mM imidazole. Fractions containing the recombinant protein were pooled, and the imidazole was removed using a PD SpinTrap™ G-25 Desalting Column (Cytiva, cat. no. 28918004). Purified proteins were stored as single use aliquots at −80 °C until further use.

### DARPin-P characterization

#### DARPin Immunoprecipitation

4.4-9.2 × 10^7^ PFU sucrose purified RSV per condition were lysed in ice cold RIPA lysis buffer (50 mM Tris pH 7.5, 150 mM NaCl, 0.1% Sodium dodecyl sulfate (SDS), 0.5% sodium deoxycholate, 1% Triton X 100) supplemented with 100 µg/ml AEBSF serin protease inhibitor (Thermo Fisher Scientific, cat. no. 78431). 12.5 µg purified DARPin-H6 was adsorbed with 10 µl Anti-DYKDDDDK Magnetic Agarose (Thermo Fisher Scientific, cat. no. A36797) for 1 h at 4°C in RIPA lysis buffer under mild rotation. DARPin-H6 adsorbed beads or untreated beads were washed in RIPA lysis buffer before incubating with virus lysate for 2 h at 4°C followed by 5 washes in ice cold RIPA lysis buffer. Proteins were eluted in 1 %SDS in D-PBS at 50°C under agitation. For proteomic analysis, 3 replicates were generated.

#### Nuclear Magnetic Resonance (NMR)

NMR experiments were carried out on a Bruker 800 MHz Avance III spectrometer equipped with a TCI cryoprobe. Protein samples were dialyzed into PBS at pH 6.4 and mixed or diluted to obtain the desired final concentration. All samples contained 7.5 % D_2_O as a lock substance. ^1^H-^15^N HSQC spectra were acquired on samples with ^15^N-P_CTD_ (50 µM) at a temperature of 288 K. The GST purification tag was removed from ^15^N-P_CTD_, which ensured that the protein was monomeric and precluded any steric hindrance that might interfere with binding. NMR data were processed with TopSpin 4.0 (Bruker) software and analyzed with CcpNMRAnalysis software^51^. Amide assignment of ^15^N-P_CTD_ was performed previously (BMRB Entry ID 26906)^42^.

#### Native agarose gel electrophoresis

Samples in the presence of 50% sucrose loading buffer were loaded on native 1% agarose gel and migration was performed in 1× Tris–Glycine buffer during 1 h 30 at 80 V before staining with amido black 10B^52^.

#### Biolayer Interferometry (BLI)

Purified MRGS(H_6_)-tagged DARPin-P (ligand) was diluted in BLI assay buffer (PBS + 0.01% bovine serum albumin (BSA) + 0.002% Tween 20, pH 7.4) at room temperature (RT). Ligands at 20 µg/ml were loaded on His1K (anti-penta-his) biosensors (Sartorius) for 150 s. Kinetic experiments were performed at 30°C with 1000 rpm shaking in 96-well black plates using the Octet Red 96e system (Fortebio). Biosensors loaded with ligands were successively equilibrated for 60 s in assay buffer (baseline step), incubated in a dilution of the analyte GST-P_CTD_ (two-fold from 350 nM to 5.5 nM) for 150 s (association step), then incubated in assay buffer for 600 s (dissociation step). One ligand-bound sensor was incubated in assay buffer as a reference to measure signal drift. As references for binding specificity, biosensors in absence of ligands were used in a parallel kinetics experiment. Real-time binding kinetics were analyzed and calculated using the Octet Red software package. Raw signal was processed using the double-reference method, by subtracting both the biosensors without ligand (unspecific signal) and the signal in the absence of analyte (drift), after baseline alignment and interstep correction at the dissociation. Kinetic modelling was done by analyzing association and dissociation signals using global fitting with a 1:1 model.

### Reporter cell line generation

#### Plasmids for lentiviral vectors

Self-inactivating (SIN) lentiviral vectors were based on the pHR vector backbone under the control of the SFFV promoter. The WPRE element was omitted to aid in obtaining low transgene expression levels (pHR-pSFFV-insert-**ΔWPRE**). The sequences of the inserted transgenes are listed in **Supplementary Table 1**.

#### Lentiviral transduction

All cell lines that stably express transgenes were generated via lentiviral transduction. Unless stated otherwise, transgenes were introduced into A549 cells. Lentivirus was produced by polyethylenimine (PEI) transfection (Polysciences Inc, cat. no. 23966) of lentiviral plasmid carrying the transgene of interest and the helper plasmids, pMD2G and psPAX2. Three days after transfection the supernatant containing lentivirus was collected and filtered (0.45 µm filter) to remove cellular debris, and was added to recipient cells of interest along with 10 mg/ml polybrene (Santa Cruz Biotechnology, cat. no. sc-134220) and subject to spin-infection for 120 min at 2000 rpm at 25°C. Following spin-infection, the medium was refreshed and after two passages, single cells from polyclonal cell populations were sorted in 96-well plates by FACS. The fluorescence expression levels for each individual cell line was carefully determined by initial investigation on the polyclone followed by screening of multiple monoclones. In general, reporter cell lines were selected with low levels of cytoplasmic fluorescence (to have minimal fluorescence background and suppress aggregate formation). Furthermore, we only selected cell lines in which RSV infection kinetics were not affected compared to parental cell lines. If a cell line required the expression of multiple transgenes, lentiviral transduction was either carried out simultaneously (maximally 2 different lentiviruses) or sequentially (minimally starting from 3 days following spin-infection). The specific cell lines used in each experiment is listed in **Supplementary Table 1**.

### RSV design, production and validation

#### Design

All viral sequences are derived from the human RSV, subgroup A, strain Long, (ATCC VR-26, GenBank accession AY911262.1). Engineered RSV strains were designed based on a previously established recombinant human RSV reverse genetics system in which unique restrictions sites have been introduced between individual RSV genes to facilitate cloning of the viral genome (reverse genetics vector pACNR-rHRSV)^16^.

The recombinant RSV virus encoding for an additional mCherry fluorescent protein (located between the P and M genes on the viral genome) encoded by the pACNR-rHRSV-mCherry was described previously^16^.

The genome expression construct encoding for recombinant upstream and downstream SunTag^eng^ RSV, were generated by designing introduction of the SunTag gene inserts at the MluI restriction site between P and M gene and BstEII restriction site between F and M2 for the upstream and downstream strains, respectively. The introduced SunTag genes were designed to contain the same gene regulatory elements, gene start (GS), gene end (GE) and 5’ and 3’ UTRs, as the N gene. Furthermore, the reporter genes were introduced such that the gene regulatory elements of the upstream and downstream genes with respect to the insert location were not disrupted. The coding sequence of the reporter genes contained a translation start codon in an optimal Kozak sequence (GCCACCATGG), followed by a sequence encoding a SunTag array and downstream gene (to generate a longer transcript allowing for more ribosomes to simultaneously translate the SunTag-encoding mRNAs, which results in higher fluorescence signal of translation sites). The upstream reporter carries a 24xSunTag-BFP and the downstream reporter carries a 12xSunTag-Kif18b. The sequences of the additional SunTag genes introduced are provided in **Supplementary Table 1**.

The engineered RSV strains carrying endogenously-tagged RSV proteins (P and L) were designed by targeting regions in the respective gene sequence that have previously been shown to be conducive for insertions with minimal disruption to viral function. An ALFA-tag was inserted into the coding sequence of P at amino acid 72^53^ to generate P-ALFA-tag^eng^ RSV and an HaloTag was inserted into the coding sequence of L at amino acid 1749^54,55^ to generate L-HaloTag^eng^ RSV. All insertions were flanked by flexible linker sequences (ALFA-Tag flanked by 5 and 3’ 6 amino acid long linkers and HaloTag flanked by a 5’ 18 amino acid and 3’ 14 amino acid linker). The engineered gene sequences are provided in **Supplementary Table 1.**

#### Generation of plasmids encoding recombinant RSV strains

The pACNR reverse genetics vector containing the RSV genomes with the upstream and downstream SunTag genes were generated by Gibson assembly and correct insertion verified by sequencing the insert and the gene upstream and downstream of the insert location.

The reverse genetics vectors carrying the P-ALFA-Tag^eng^ and L-HaloTag^eng^ RSV strains were generated by transformation-associated recombination (TAR) cloning in yeast as described previously^56^. Briefly, overlapping PCR products containing the RSV genome, ALFA-tag/HaloTag, and TAR vector (pCC1BAC-His3)^57^ were transformed into yeast strain VL6-48N^58^. Yeast-assembled full-length clones were isolated, amplified in *E. coli* strain EPI300, and sequences were verified by full-plasmid nanopore sequencing (Plasmidsaurus, USA), where remaining uncertainties in the viral sequence were confirmed by Sanger sequencing.

#### RSV rescue and concentration

Recombinant WT and tagged RSV strains were rescued by reverse genetics and amplified in HEp-2 cells as previously described^59^. In brief, the appropriate pACNR reverse genetics vector (1.25 µg) was co-transfected together with the expression vectors pCITE-N (1 µg), pCITE-P (1 µg), pCITE-L (0.5 µg) and pCITE-M2-1 (0.25 µg)^16^, each under the control of the T7 promoter^54^ in BSR T7/5 using Lipofectamine3000 (Thermo Fisher Scientific, cat. no. L3000001), according to the manufacturer’s protocol. 3 days following transfection, virus was collected and efficiency of rescue assessed by plaque assay carried out on HEp-2 cells. The viruses were amplified for a subsequent 3 to 4 passages on HEp-2 cells at an MOI of 0.01 PFU/cell to minimize formation of defective viral genomes. For the P-ALFA-tag^eng^ and L-HaloTag^eng^ RSV strains, the rescue protocol was similar, with the exception that the helper plasmids were in pcDNA3 (BEI Resources, cat. no. NR-36462, NR-36463, NR-36464, NR-36461), transfection of BSR T7/5 cells was performed using Lipofectamine LTX (Thermo Fisher Scientific, cat. no. A12621), passaging was performed on Vero cells, and viruses were titrated by 50% tissue culture infective dose (TCID_50_) assay. The final stocks were grown on Hep-2 cells.

Viral supernatants from passage 3-4 were precleared for cells and debris by centrifugation and subsequently concentrated. Concentration was done either via polyethylene glycol (PEG)-mediated precipitation (viral supernatant was stirred with 10% PEG6000 (50 mM Tris-HCL, pH 7.4, 150 mM NaCl, 1 mM EDTA,) for 3 h at 4°C, followed by centrifugation at 3,500 RCF for 30 min at 4°C), or by pelleting through a 10% sucrose-containing buffer (50 mM Tris-HCl, pH 7.4, 100 mM NaCl, 0.5 mM EDTA) (centrifuge viral supernatant in 10% sucrose buffer (4:1 v/v ratio) at 10,000 RCF for 4 h at 4°C). The virus containing pellets were resuspended in 10% sucrose in PBS, snap-frozen and stored at −80°C as single-use aliquots.

#### Viral growth assays

Viral growth assays were performed to assess the fitness of the engineered RSV strains with respect to their WT counterpart.

The upstream and downstream SunTag^eng^ RSV strains growth kinetics were assessed on A549 cells. A549 cells were inoculated with virus at an MOI of 3 PFU/cell (in DMEM + 10% FBS + 1% Pen-Strep) for 1 h, following which cells were washed in PBS and supplemented with fresh media (DMEM + 10% FBS + 1% Pen-Strep). New progeny virus production was assessed up to 5 days post infection by collecting samples at multiple time points along the time course. The viral titers of the samples were determined by plaque assays on HEp-2 cells.

The growth kinetics of the P-ALFA-tag^eng^ and L-HaloTag^eng^ RSV strains were assessed on Vero cells via qPCR. Cells were infected with an MOI of 0.04 PFU/cell and supernatant was collected up to 48 hours post infection in MagNA Pure External Lysis Buffer (Roche, cat. no. 06374913001) for RNA isolation. 120 μL sample was mixed with 50 μL AMPure XP Reagent (Beckman Coulter, cat. no. A63880), loaded in a 96-wells PCR plate, washed three times with 70% ethanol using a DynaMag-96 Side Skirted Magnet and RNA was eluted in 50 μL RNase-free H_2_O. RNA, 5 μL per 20 μL reaction volume, was reverse-transcribed and amplified in one reaction using the TaqMan Fast Virus 1-step Master Mix (Thermo Fisher Scientific, cat. no. 4444432) according to manufacturer’s protocol on a StepOnePlus System. RSV-A primers with a FAM-probe were used (**Supplementary Table 1**) and a serially diluted RSV-A plasmid was included as a reference.

### Cell culture and RSV infection for imaging-based studies

#### Inoculum for RSV infection

One day prior to imaging, the required A549 cell lines were seeded on a 96-well glass-bottom plates (Brooks Life Science Systems, cat. no. MGB096-1-2-LG-L and Ibidi, cat. no. 89627) such that cells were at ∼80-90% confluency at the point of viral inoculum addition. For fixed cell experiments, the viral inoculum was made in cell culture medium (DMEM + 10% FBS + 1% Pen-Strep). For live-imaging experiments viral inoculum was made in imaging medium (CO_2_-independent Leibovitz’s-L-15 media (Thermo Fisher Scientific, cat. no. 21083027) supplemented with 10% FBS and 1% Pen-Strep).

For live cell assessment of RSV infection progression and success, a 1:1000 dilution of RSV Glycoprotein G Antibody (133) (referred to as AbG-fluoro) (NOVUS Biologicals, Janelia Fluor 646: cat. no. NBP2-50411JF646 and Alexa Fluor 405: cat. no. NBP2-50411AF405) was added directly into the viral inoculum. For visualization of the L-HaloTag^eng^ RSV the viral inoculum was supplemented with 5 nM HaloTag Ligand JFX650^60^. Both fluorescent media supplementations gave negligible fluorescence background and was maintained on cells until experimental end point. The details of viral inoculums used in each experiment is listed in **Supplementary Table 1**.

#### Multiplicity of infection (MOI) used for imaging-based studies

For imaging studies MOI was based on the fraction of cells containing vRNPs at 6h following viral inoculum administration. In fixed cell experiments with WT A549 cells this was assessed by the presence of N^+^/F^−^ foci (antibody staining) and in live cell experiments this was assessed in the P^exo^-fluoro transgenic cell line by the presence of P^exo^ foci. The MOI value established using this imaging-based method is 3 orders of magnitude higher than the MOI calculated using plaque assay-based viral titration on HEp-2 cells (0.1 imaging-based MOI relates to 0.0001 PFU/cell). Unless otherwise specified, the imaging-based MOI used was between 0.05-0.25.

#### Chemical inhibitors

The following inhibitors were used in this study: PC786 (15 µM, RSV polymerase inhibitor) (MedChemExpress, cat. no. HY-102038), TMC353121 (180 nM, RSV fusion inhibitor) (MedChemExpress, cat. no. HY-11097), emetine dihydrochloride (referred to as emetine, 50 µg/ml, inhibits translation by preventing translocation of the tRNA-mRNA complex) (Merck, cat. no. 324693) and Puromycin dihydrochloride (referred to as puromycin, 0.1 mg/ml, a tyrosyl-tRNA mimic that blocks translation by releasing elongating polypeptide chains) (Thermo Fisher Scientific, cat. no. A1113803). When used, PC786 and emetine were added directly with the viral inoculum, and TMC353121 and puromycin were added to infected cells at indicated time points. Following addition, the inhibitors were maintained on cells until experimental end point. For multi-day inhibitor treatment protocols, the inhibitors were refreshed after each 24 h.

#### Assessing infecting-vRNPs

Assessment of the mobility, viral protein and RNA levels on infecting RSV vRNPs were carried out in the presence of the translation inhibitor, emetine. Emetine containing inoculum was maintained on cells for 1 h, following which the RSV fusion inhibitor, TMC353121 was added to the inoculum to prevent subsequent infections. 3 h post fusion inhibitor addition the cells were assayed, either by live imaging to assess foci mobility or fixed and subject to antibody or smFISH staining protocols. This time point was chosen to ensure that all infecting vRNPs had fully separated such that infecting individual vRNPs could be assessed. For vRNP mobility assessment a higher MOI (MOI 2) was used, such that cells simultaneously infected with both DARPin-P^+^ and DARPin-P^−^ vRNP-containing virions could be identified, and the kinetics of both vRNP states assessed in the same cell. This approach was adopted to exclude the cell to cell heterogeneity impacting mobility measurements.

### Immunofluorescence

#### Antibody conjugation

RSV Nucleoprotein Antibody (RSV3132 (B023)) (referred to as anti-RSV N) (NOVUS Biologicals, cat. no. NB100-64752) was fluorescently-labelled using N-hydroxysuccinimide (NHS)-esters (Alexa Fluor™ 555 NHS ester and Alexa Fluor™ 647 NHS ester) (Thermo Fisher Scientific, AF555: cat. no. A20009, AF647: cat. no. A20106) for use in immunofluorescence. The antibody was buffer-exchanged and concentrated with a 10 kDa MWCO PES concentrator (Thermo Fisher Scientific, cat. no. 88535) to yield 3 mg/ml antibody solution in 7.5% sodium bicarbonate buffer, pH 8.3 (Thermo Fisher Scientific, cat. no. 25080094). The dye-NHS stocks were prepared in anhydrous DMSO (Thermo Fisher Scientific, cat. no. AM9342). A 3-fold molar excess of the dye-NHS were incubated with 1 mg of the antibody for 2 h at RT with continuous rotation. Free dye was removed using a PD SpinTrap™ G-25 Desalting Column (Cytiva). Protein concentration and degree of labelling (DOL, fluoro-to-protein (F/P) ratio) were determined using NanoDrop spectrophotometric absorbance values. All antibody conjugations yielded a DOL of 2-3 dyes per antibody.

#### Immunofluorescence of infected cells and RSV virions on glass

Infected cells and uninfected control cells were fixed with 4% paraformaldehyde (Thermo Fisher Scientific, cat. no. 043368.9M) for 10 min at RT at indicated time points. All fluorescence of genetically encoded fluorescent proteins is maintained following this fixation and antibody staining protocol. If virions attached to the extracellular membrane of cells needs to be stain, RSV Glycoprotein F Antibody (11-2-F3) (referred to as anti-RSV F) (NOVUS Biologicals, DyLight550: cat. no. NBP2-50412R, Alexa Fluor 488: cat. no. NBP2-50412AF488) staining is carried out prior to permeabilization and blocking. Antibody diluted to the appropriate concentration in PBS + 10% FBS was added to cells and incubated for 1 h at RT (1:500 anti-RSV F (AF488/DL550)). Following incubation samples were washed three times in PBS. Samples were subsequently blocked and permeabilized, simultaneously, with blocking buffer (PBS + 10% FBS + 0.05% Triton-X) for 1 h at RT. Antibodies and recombinant proteins were diluted to the appropriate concentration in blocking buffer and were incubated for 1 h at RT (recombinant proteins: 1:1000 of 0.4 mg/ml DARPin-P-mRuby3; antibodies: 1:1000 of 0.5 mg/ml anti-RSV N (AF555/AF647), 1:1000 RSV Phosphoprotein Antibody (2H102) [Janelia Fluor® 646] (referred to as anti-RSV P (JF646)) (NOVUS Biologicals, cat. no. NBP2-50276JF646), 1:1000 RSV M2-1 Protein Antibody (5H5) [Alexa Fluor® 647] (referred to as anti-RSV M2-1 (AF647)) (NOVUS Biologicals, cat. no. NBP2-50481AF647)). Following incubation, samples were washed three times in wash buffer (PBS + 0.05% Triton-X). If whole cell segmentation was required, total protein was stained using Pacific Blue succinimidyl ester (Thermo Fisher Scientific) (1:2000 of 1 mg/ml stock solution made up in anhydrous DMSO) in PBS for 10 min at RT followed by three washes in wash buffer. Samples were kept in PBS at 4°C until imaging.

To visualize extracellular G protein expression on RSV infected cells, 1:1000 dilution of AbG-fluoro (JF646/JF405) was added directly to the cell culture media and maintained for 30 min at 37°C. Following incubation, the cells were washed three times in PBS and fixed as described above.

Virion protein levels were assessed by incubating RSV inoculum in PBS (total volume 100 µl) for 1 h at 37°C directly in a well of a 96 Well Square Glass Bottom plate. Following incubation, virions were fixed by adding 4% paraformaldehyde directly into the inoculum and incubating for 15 min at RT. The antibody staining was carried out as described above. In addition to the antibodies listed above, 1:1000 FluoTag®-X2 anti-ALFA [Atto 488] (referred to as anti-ALFA (Atto488)) (NanoTag Biotechnologies, cat. no. N1502) was used for virion labeling.

All antibodies, purified proteins and staining protocols used in this study have been optimized to yield minimal to no background foci signal, such that the staining can be used to evaluate RSV infections at a single vRNP resolution. The specific antibody-fluorescent conjugations used per experiment is listed in **Supplementary Table 1**.

### Single-molecule fluorescence in situ hybridization (smFISH)

#### smFISH probe generation

Stellaris probe designer (https://www.biosearchtech.com/support/tools/design-software/stellaris-probe-designer) was used to design probes targeting the entirety of the negative-sense RSV genome, intergenic regions of the positive-sense RSV antigenome (to minimize cross reactivity with viral mRNAs), positive-sense RSV transcript mRNAs and SunTag mRNA. Oligonucleotide probes were ordered from Integrated DNA technologies (sequences are listed in **Supplementary Table 1**). All probes targeting a single RNA were pooled and labelled with ddUTP-coupled ATTO 565 (ATTO-Tec, cat. no. AD565-31) or ATTO 633 (ATTO-Tec, cat. no. AD633-31) dyes using terminal deoxynucleotidyl transferase (Thermo Fisher Scientific, cat. no. EP0161) as previously described^61^. For the RSV all mRNA probe set probes targeting all RSV transcripts were pooled. Fluorescent probes were purified by ethanol precipitation and resuspended in nuclease-free water to a final stock concentration of 30 µM.

#### smFISH probe hybridization

smFISH staining procedure was carried out as previously described ^62^. Briefly, following fixation, cells were washed 3 times in PBS, permeabilized in 100% ethanol for 1 h at 4°C and then washed 3 times for 15 min in smFISH wash buffer (2xSSC, 10% formamide (Thermo Fisher Scientific, cat. no. 17899) in nuclease free water) at RT. Labelled smFISH probes were diluted to 10 nM in hybridization buffer (1% dextran sulfate (Merck, cat. no. D8906), 2xSSC, 10% formamide, 1 mg/ml tRNA (Merck, cat. no. R1753), 2 mM Ribonucleoside vanadyl complex (referred to as RVC, New England Biolabs, cat. no. S1402S), 200 µg/ml BSA (Thermo Fisher Scientific, cat. no. AM2616) in nuclease free water) and hybridization was performed at 37°C for 16 h. Unbound smFISH probes were removed by 3x 30 min washes in smFISH wash buffer at 37°C and a 15 min wash at RT. If whole cell segmentation was required, total protein was stained using Pacific Blue succinimidyl ester (1:2000 of 1 mg/ml stock solution) (Thermo Fisher Scientific, cat. no. P10163) in PBS for 10 min at RT followed by a 15 min wash with smFISH wash buffer at RT. Samples were stored at 4°C and imaged in smFISH imaging buffer (10 mM Tris pH 8, 2xSSC, 0.4% glucose, supplemented with glucose oxidase (Merck, cat. no. G2133) and catalase (Merck, cat. no. C3515)). Imaging was performed within 3 days after probe hybridization. The specific smFISH probe sets used in each experiment is listed in **Supplementary Table 1**.

#### Combining smFISH and N antibody staining

To combine smFISH with anti-RSV N antibody staining, the wash buffer was additionally supplemented with 3% BSA and the fluorescently-conjugated antibody was directly added into the smFISH hybridization buffer at a 1:100 dilution. The remaining protocol is carried out as detailed above.

#### Combined smFISH and STAb staining

To combine smFISH with STAb-sfGFP staining, after the three washes with smFISH wash buffer at 37°C, samples were incubated with purified scFv-sfGFP-STrepII^15^ (referred to as STAb-sfGFP, 1:100 dilution in PBS + 10% BSA) for 1 h at RT. Following incubation, samples were washed three times in PBS. A final RT wash step of 30 min in smFISH wash buffer was then performed prior to sample storage and imaging in smFISH imaging buffer.

#### Combining smFISH staining with fluorescence originating from transgenic cell lines

For the P^exo^-fluoro cell line, used to visualize all RSV vRNPs, the foci signal (in the AausFP1 fluorescent protein) is well retained following the smFISH protocol and can be visualized simultaneously with the smFISH foci. For the DARPin-P visualizing transgenic cell line, the foci signal (in the BFP fluorescent protein) is somewhat compromised following the smFISH protocol and as such needs to be first imaged post fixation, but prior to ethanol permeabilization, and then followed by the smFISH protocol. When DARPin-P signal visualization with smFISH signals were required, experiments were only carried out in the P^exo^-fluoro and DARPin-P-fluoro combined transgenic line. Acquiring the P^exo^ signal both in the post fixation and post smFISH imaging rounds allowed it to be used to align the data from the two acquisition rounds, ensuring that accurate values for the DARPin-P signal (from post fixation imaging) could be used for the cells being analyzed.

### Flow cytometry-based assessment of successful RSV infections

A flow cytometry -based assay that uses the expression of viral G protein by infected cells was used as a read-out of successful RSV infections. This assay was used to assess RSV infection success in the DARPin-P-fluoro transgenic A549 cell line compared to WT A549 cells. WT A549 cells and DARPin-P-fluoro cells were plated, in parallel, on a 96-well glass-bottom imaging plate (for establishment of MOI) and a 24-well plate (for flow cytometry assessment). The next day cells were infected with RSV (0.001 PFU/cell). *MOI establishment*: Following 6 h of infection, the cells plated on the imaging plate were fixed and subject to antibody staining for anti-RSV N and F and the number of infected cells (N^+^/F^−^) determined to obtain a MOI. *Flow cytometry assessment*: At 6 h cells infected for flow cytometry assessment were treated with an RSV fusion inhibitor (TMC353121) to prevent subsequent infections. Flow cytometry assessment was carried out at 24 and 48 h following viral inoculum addition. An hour prior to flow cytometry, infected cells (and uninfected control cells) were incubated with AbG-fluoro (1:1000, in cell culture media) for 45 min at 37°C. After antibody incubation cells were rinsed in PBS, trypsinized, fixed (in suspension, with 4% paraformaldehyde for 10 min at RT with constant rotation) and suspended in PBS. Flow cytometry analysis was performed using Cytoflex S (Beckman Coulter) and CytExpert software (see **Supplementary Fig. 1**). The percentage of successful infections in each cell line was calculated in relation to the fraction of infected cells at 6 h post inoculum addition (MOI) to determine whether expression of the DARPin-P-fluoro reduced the efficiency of RSV infection success.

### Microscopy

#### Microscope

All fluorescence microscopy was carried out on a Nikon TI2 inverted microscope equipped with a Yokagawa CSU-X1 spinning disc and a Prime 95B sCMOS camera (Photometrics). Unless state otherwise, imaging was performed using a x60/1.40 NA oil-immersion objective. Image acquisition was performed using NIS Elements software, making use of the ‘perfect focus system’ to correct for Z drift during time-lapse imaging experiments. The microscope was equipped with a temperature-controlled incubator and imaging for live-cell experiments was performed at 37°C and for fixed-cell experiments at RT.

#### Image acquisition

For time-lapse imaging experiments, x, y positions for imaging were randomly selected. Position selection, was carried out immediately following viral inoculum addition and prior to observing foci in infected cells. Images were acquired every 5, 10 or 20 min (exact intervals are recorded in the figure legends) for 6 h (short-term time lapse imaging) or 12-48 h (for over-night and longer-term time lapse imaging), using 50-70 ms exposure times (50 ms for the BFP channel and 70 ms for all other channels). Multiple Z-slices (∼9-15 slices with a 0.8 µm step size) were imaged for each channel ensuring the entirety of the cell was captured.

For experiments in which the diffusion speed of infecting vRNPs was assessed, cells expressing P^exo^-fluoro and DARPin-P-fluoro were infected in the presence of a translation inhibitor (emetine) for 4 h (see section, ‘Assessing infecting-vRNPs’ for details). Infected cells were selected based on the presence of DARPin-P^+^ and DARPin-P^−^ vRNPs. A single Z-slice for both the P^exo^-fluoro and DARPin-P-fluoro channels were acquired in the region of the cell that most optimally visualizes the foci, using 15% laser power with 50 msec exposure (for both channels) at a 1 sec interval for 1 min.

For fixed cells analysis of immunofluorescence or smFISH staining, x, y positions for imaging were selected either randomly using the cell segmentation channel (when data regarding the frequency of a phenotype was analyzed) or targeted using the channel of interest (when the frequency of a phenotype is not assessed). For smFISH staining where the transgenic cell line DARPin-P-fluoro signal is acquired post-fixation, the same x, y positions are used to acquire the smFISH staining. Images were acquired at 45% laser power and 70 ms exposure (for all lasers) with a 0.8 µm step size and a Z-stack slice number that ensures the entirety of the cell is captured.

#### Post-acquisition data processing and move generation

Maximal intensity projections for all Z-slices (where multiple Z-slices were acquired) were generated using NIS Elements software (Nikon) and all downstream analyses were performed on these projections.

The single-object-grid tool (https://github.com/TanenbaumLab/SingleObjectGrid) is used to crop a small field of view around tracked cells in order to create a stabilized timeseries with the cell centroid centered. The napari-animation plugin is used to generate video output from the napari viewer to create supplemental videos (https://napari.org/napari-animation/).

### Quantification

#### Foci quantification

For all analyses infected cells were manually identified by the presence of RSV vRNPs (as determined by the P^exo^ foci, anti-RSV N antibody staining or (−)vRNA smFISH staining). Only cells that were completely in the FOV were included in the analysis.

For fixed cell analysis infected cells were manually segmented in ImageJ and foci detection and foci integrated intensity quantification were performed using the ‘detect particles and colocalization analysis’ function in the ComDet v.0.5.4 plugin in ImageJ (https://github.com/UU-cellbiology/ComDet). Plugin spot detection parameters were optimized for each type of foci analyzed. The intensity thresholds used in the analyses were established for each individual experiment using the uninfected control samples. The specific parameters used were: smFISH viral N mRNA spot detection; approximate particle size of 3 µm, intensity threshold range ≥15-30 (i.e. ≥15-30 times the signal-to-noise-ratio), large particles not included; smFISH viral genome and antigenome spot detection; approximate particle size of 4 µm, intensity threshold range ≥5-50, large particles included; transgenic cell line- and antibody staining-derived RSV vRNP spot detection; approximate particle size of 4 µm, intensity threshold range ≥5-200, large particles included. For individual vRNP spot detection (including smFISH, transgenic cell line and antibody staining-derived foci) an additional filtering step was introduced, whereby only spots that had an N Area value >6 and <30 (pixels in foci) were included in the analysis. Intensity-based cutoffs for each channel were used to call colocalization. These cutoff values were established for each individual experiment using both the uninfected control sample and individual channel of interest. The lowest intensity where a foci was detected (individual channel of interest) above background was used as the intensity cutoff to call positive and negative values. Foci integrated intensity values (sum of all pixels intensities inside the segmented spot area minus average spot-specific backgrounds. Background value is calculated as the average intensity of pixels along the perimeter of a bounding rectangle surrounding the segmented spot) <0 were converted to 0 to aid with analysis and data visualization. Integrated intensity values were normalized (to the median value of DARPin-P^−^ foci, or to the maximum intensity value) to allow for optimal data comparison. For antibody staining experiments, only genomes that were N^+^/F^−^ were considered to be inside cells. If the P^exo^-fluoro transgenic cell line was utilized, only P^exo+^ foci were considered to be vRNPs inside infected cells.

Quantification of the time foci appeared and the number of foci present at each time frame in live-cell time lapse imaging movies was carried out manually. Only cells that remained completely within the FOV for the entire duration of the analysis were included. For experiments where infection was followed from the moment of viral cell entry, only cells where a minimum of 2-time frames prior to viral entry was observed were included. For vRNP analysis, only foci that were present for a minimal of 2 consecutive time frames were considered in the analysis to minimize false foci calling. For assessing the DARPin-P status, infections were classified as DARPin-P positive if one or more of the infecting vRNPs were DARPin-P^+^ even if the cell additionally had DARPin-P^−^ vRNPs. Quantification of the number of translating viral reporter mRNAs per cell (infection with the upstream and downstream SunTag^eng^ RSV strains) was performed, based on previously described guidelines ^15^. In brief, the number of translation sites at each individual time frame were assessed until individual translation sites could no longer be detected (high translation rate leading to the depletion of free cellular STAb due to the cytoplasmic accumulation of mature SunTag peptides)^63^. AbG-fluoro signal accumulation was also assessed manually and recorded at the first time frame positivity could be observed. Foci calling (P^exo^ and DARPin-P signal for vRNPs and translation sites for the the SunTag^eng^ RSV downstream engineered strain) and AbG-fluoro accumulation calling was independently validated by another lab member. If intensity quantification of foci in time-lapse movies was performed, the cell of interest was manually segmented at each time point and foci intensity quantified using the ComDet plugin. To facilitate visual inspection of general trends in heterogenous intensity values over time (for individually tracked foci), moving averages are plotted with a sliding window of 3 time points.

#### Quantification of viral transcription rate

Since the number of translating SunTag mRNAs closely correlated with the number SunTag mRNAs (**Extended Data Fig. 5f**), the increase in the number of translating SunTag mRNAs over time was used as a proxy to calculate viral transcription rate. The number of viral mRNA translation sites within an 1 h period since initial observation was plotted against time. Graphs for individual infections were fit with a linear function using GraphPad prism (v 10.1.0) and the slope of this function is used as the viral transcription rate.

#### Calculating the mobility of RSV vRNPs

vRNP foci detection and tracking was carried out using an automated inhouse optimized image analysis pipeline that included an illumination correction, spot detection and spot tracking module. *Illumination correction*: Images for illumination correction are acquired using a 100 mg/ml solution of fluorescein disodium salt in water ^64^. In total 147 exposures at different positions are recorded and median intensity projected to produce a flat field image. Dark field images are generated in an equivalent manner but with excitation light off. To transform the flat field image into the final correction matrix the dark field image is subtracted and the result is divided by the mean intensity of the full image. To perform flat field correction on raw images, first the dark field image is subtracted, followed by division by the correction matrix. *Spot detection*: Image data analysis is performed using Python (3.9.7). ND2 imaging files are loaded using the nd2 github.com/tlambert03/nd2 (0.5.3) library.

Fluorescent foci are detected using Laplacian of Gaussian detection from the blob_log function of scikit-image (0.22.0)^65^ with settings thresholdLoG = 10**-3, min_sigma = 2, max_sigma = 6, overlap = .75. The returned sigma values per detection are used to classify *’normal’ and ‘large’ IBs* at a threshold of 2.5. *Spot tracking*: Detected spots are tracked over time using the LAPTrack library^66^ with squared euclidian distance cost cut-off 25, track splitting and merging disabled, maximum frame gap of 3 frames. Data exploration and annotation is performed with the interactive image visualization suite Napari^67^ (0.4.18). DARPin-P state is manually assigned after rearranging and stabilizing all tracked foci into a grid using the single-object-grid tool (https://github.com/TanenbaumLab/SingleObjectGrid). Only tracks with a minimum length of 10 frames are used for further analysis. Diffusion coefficient of each track was calculated by fitting the first 5 time-points of the mean square displacement (MSD) curve using inhouse MATLAB (The Mathworks, Inc.) scripts.

#### Proteomic quantitative analysis

Sample preparation was done via the bead-based single-pot, solid-phase-enhanced sample-preparation (SP3) and analyzed by LC/MS/MS. Protein identification and quantification were performed using Andromeda search engine implemented in MaxQuant under default parameters. Peptides were searched against reference Uniport datasets and RSV custom proteome derived from HRSV subgroup A, strain Long (GenBank accession: AY911262). False discovery rate (FDR) was set at 1% for both peptide and protein identification. MaxQuant search was performed with “match between run” activated in the searches. MaxQuant (proteinGroups) outputs were used for downstream relative quantification analysis. Proteins highlighted as potential contaminant and reverse were filtered out, together with proteins with missing values for more than 4 of the 6 conditions. R-package limma (3.52.4)^68^ was used for statistical analysis for protein intensities, applying a moderated t-test with p-values adjusted for multiple testing using Benjamini-Hochberg methodology. Data was visualizes using ggplot2 (3.4.4).

### Statistical analysis

All statistical analyses performed using GraphPad prism (v 10.1.0) (GraphPad Software). Unless stated otherwise, statistical tests were performed using a *P* value of 0.05 as a cut-off for significance and assuming normal distribution of experimentally determined averages. The type of test and the type of error bars used in figures are indicated in the figure legend or in the figure itself. The *n* values are recorded in the figures as well as included in **Supplementary Table 1**. Additionally **Supplementary Table 1** presents an overview of the number of experimental repeats, the total number and type of observations per condition, as well as situations where the same raw data has been used for multiple independent analyses. Graph creation was also performed in GraphPad.

### Data availability

Further information and requests for resources and reagents should be directed to and will be fulfilled by the lead contact, Marvin E. Tanenbaum. The unique/stable reagents generated in this study are available from the lead contact with a completed Material Transfer Agreement. The plasmids generated in this study for the vRNP visualization tools (P^exo^-fluoro and DARPin-P-fluoro) will be deposited in Addgene (*at publication*). A selection of raw imaging data has been made available through figshare.

### Code availability

No custom code was used to generate or process the data described in this manuscript.

## Acknowledgments

We thank Y. Demyanenko and S. Mohammed for performing mass-spectrometry analysis, S. Yang for providing script and support for MSD calculation, B. Verhagen for help with cell line establishment, J. Schokolowski for help with independent validation of time-lapse analysis, H.J. Hamstra and A. Gelderloos for plasmid generation and primary rescue of the P-ALFA-Tag and L-HaloTag RSV strains, M. Müller for discussions on the viral labeling strategy, L. Lavis for the Halo Tag ligands, S. Vashee for providing the TAR vector, N. Kouprina for providing the yeast strain, V. Thiel for help with the TAR cloning protocol, L. Bont lab for help with the virus concentration protocol and Hubrecht Institute flow cytometry core facility for facilitating cell sorting. We also thank members of the Tanenbaum lab, Bont lab and L. Meyaard for discussions and J. Schuijers for comments on the manuscript. This work was financially supported by an ERC CoG grant to M.E.T (ERC, VirIm,101044794), the Howard Hughes Medical Institute through an international research scholar grant to M.E.T. (HHMI/IRS 55008747), an EMBO Postdoctoral Fellowship to D.R. (ALTF 102-2021), D.R., S.B., M.B., R.B., and M.E.T. were supported by the Oncode Institute, which is partly funded by the Dutch Cancer Society (KWF), A.J.L. and P.B.v.K were financially supported by the Dutch Ministry of Health, Welfare and Sport. A.C. is funded by the European Research Council (ERC) Consolidator Grant ‘vRNP-capture’ 101001634 and the Medical Research Council (MRC) grants MR/R021562/1 and MC_UU_00034/2. While this work is funded by the European Union, views and opinions expressed are however those of the author(s) only and do not necessarily reflect those of the European Union or the European Research Council Executive Agency. Neither the European Union nor the granting authority can be held responsible for them.

## Author contributions

Conceptualization, D.R., S.B. and M.E.T.; Methodology, D.R., S.B. and M.E.T.; Investigation, D.R., M.G., S.B., C.S., J.S., A.J.L., R.B., M.N. B.D. and S.F.; Formal analysis, D.R., M.G., S.B., C.S., J.S., A.J.L., M.B., and M.N.; Validation, D.R., and S.B.; Visualization, D.R., M.G., S.B., C.S., J.S., A.J.L, M.B., and M.N.; Resources, M.-A.R.-W., B.D., S.F., A.P., M.G., J.-F.E., and P.B.v.K.; Writing-original draft, D.R., and M.E.T; Writing-review & editing, all authors; Supervision, M.G., J.-F.E., P.B.v.K., A.C., A.P. and M.E.T.; Funding acquisition, D.R., A.P., J.-F.E., C.S., P.B.v.K., A.C., and M.E.T.

---

**Extended Data Fig 1.**
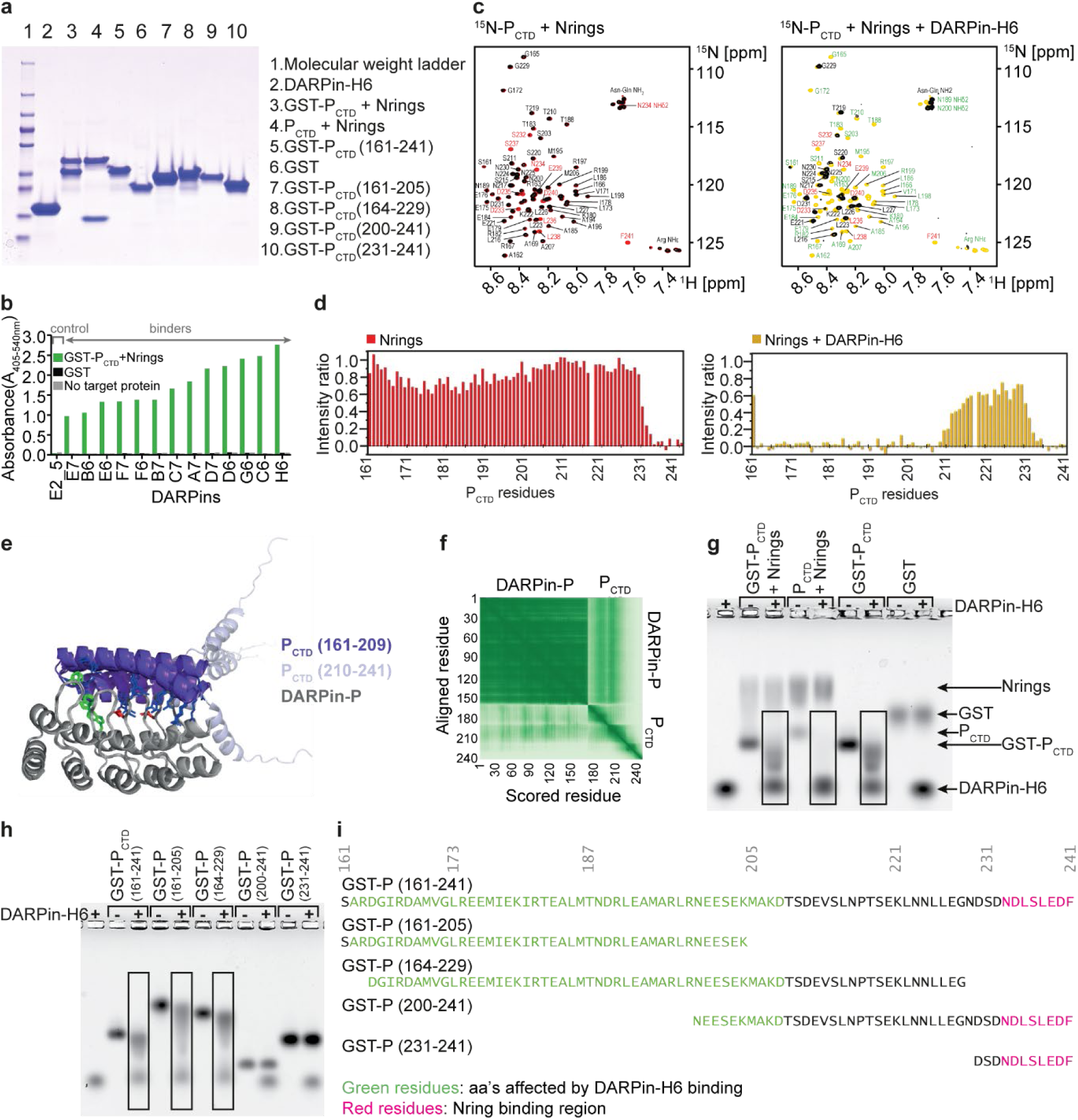
Identification and *in vitro* characterization of a DARPin that binds RSV phosphoprotein. **a,** SDS-PAGE and Coomassie blue staining of purified DARPin-H6, P_CTD_+Nring protein complex and tags used as a target for DARPin selection and P_CTD_ fragments. The first lane shows the molecular weight ladder. **b**, DARPins that bind RSV vRNPs were identified via ribosome display. ELISA measurements show specific binding of different DARPins to the target protein complex. The absorbance for 13 target binding DARPins and a non-binding control DARPin (DARPin E2_5)^44^ (green bars) is shown. GST (black bars) and no target (grey bars) were used as controls. DARPin-H6 was selected for further characterization. **c, d,** 2D NMR spectra of RSV ^15^N-P_CTD_ in the presence of DARPin-H6 and Nrings. (**c**) ^1^H-^15^N HSQC spectra of 50 µM ^15^N-P_CTD_ were measured after addition of 1 molar equivalent of RSV Nrings, and after addition of 1 molar equivalent of each RSV Nrings and DARPin-H6 (both spectra in black). The Nring spectrum (in red) and Nring+DARPin-H6 spectrum (yellow) were superimposed on the spectrum of ^15^N-P_CTD_ alone. Assignments of signals that were fully broadened out due to addition of Nrings and DARPin-H6 are indicated in red and green, respectively. Other assignments are in black. (**d**) Backbone amide signal intensities in ^1^H-^15^N HSQC spectra of ^15^N-P_CTD_ were measured for each residue, except proline 218, in the presence of 1 molar equivalent of RSV Nrings, or both RSV Nrings and DARPin-H6, as well as for ^15^N-P_CTD_ alone. The bar diagrams represent the intensities measured in the presence of interaction partners divided by the intensities of the reference spectrum (intensity ratio). Addition of Nrings to P_CTD_ resulted in broadening of the backbone amide signals corresponding to the last 10 residues at the C-terminal end of the P_CTD_, amino acids that were previously identified as an Nring binding region^42,69^. When RSV Nrings were added to the DARPin-H6−P_CTD_ complex, the spectral perturbations observed for the ternary mixture were the sum of individual perturbations induced by DARPin-H6 and Nrings (see also Fig. 1b, c), indicating that DARPin-H6 binds to P_CTD_ alone and P_CTD_ in complex with Nrings in a similar manner. **e, f,** AlphaFold3^70^ prediction of the DARPin-H6:RSV P_CTD_ complex (https://alphafoldserver.com/, accessed on December 5, 2024). The 40 residue P_CTD_ region identified as the DARPin-H6 binding region is larger compared to the size of DARPin-H6. DARPin-H6 would span 65 Å if a single extended α-helix was formed. This is twice the distance between the first and the fourth loop of the variable region of DARPin-H6, and suggests that the P_CTD_ region takes on a compact, folded conformation in the P_CTD_+DARPin-H6 complex. The NMR data strongly suggests that tumbling of the P_CTD_ (aa 162-209) region is slowed down due to binding DARPin-H6, implying that this region of the P_CTD_ is indeed folded or structured in the DARPin-H6−P_CTD_ complex. Previous studies have identified two transient α-helices, α_C1_ and α_C2_, in the largely unstructured CTD of RSV P by NMR^42^, and the P_CTD_ has been shown to also adopt a structured conformation when in complex with L^29^, consistent with our data that this region can adopt a structured conformation. AlphaFold3 (AF3) was used to predict the potential structure of the DARPin-H6−P_CTD_ complex. The protein query sequences were DARPin-P and P_CTD_ (aa 161-241). (**e**) Superimposition of the 4 best ranked complex models, with DARPin-P in grey, P_CTD_ aa 161-209 in purple, and P_CTD_ aa 210-241 residues in light blue. Protein structures were visualized with Pymol (Schrodinger, 2010, https://pymol.org/). P arginine side chains are shown in blue sticks. DARPin-H6 residues that interact with P arginines are in sticks (acidic residues - red, aromatic residues - green). The models generated predicted that the 50 N-terminal residues of the P_CTD_ formed two parallel helices that interact with the variable region of DARPin-H6. (**f**) Predicted aligned error (PAE) matrix of the AlphaFold3 prediction (color code for expected position error: 0 to 30 Å from dark green to white). **g,** DARPin-H6 interacts with RSV P, both in its free and Nring bound form. DARPin-H6 was co-incubated with the P_CTD_+Nrings target protein complex or its individual components (including tags) and interactions were assessed by band shift on native agarose gel. Boxes indicate lanes where interactions are observed. **h, i,** NMR-identified DARPin-H6 interaction region on RSV P was further validated by assessing complex formation of DARPin-H6 with fragments of the P_CTD_. (**h**) Complex formation was visualized by band shift on native agarose gel. Boxes indicate lanes showing complexes. (**i**) The amino acid sequence of the P_CTD_ and its fragments are shown. P_CTD_ residues identified to interact with DARPin-H6 by NMR are in green and Nring binding residues are in magenta.

**Extended Data Fig 2.**
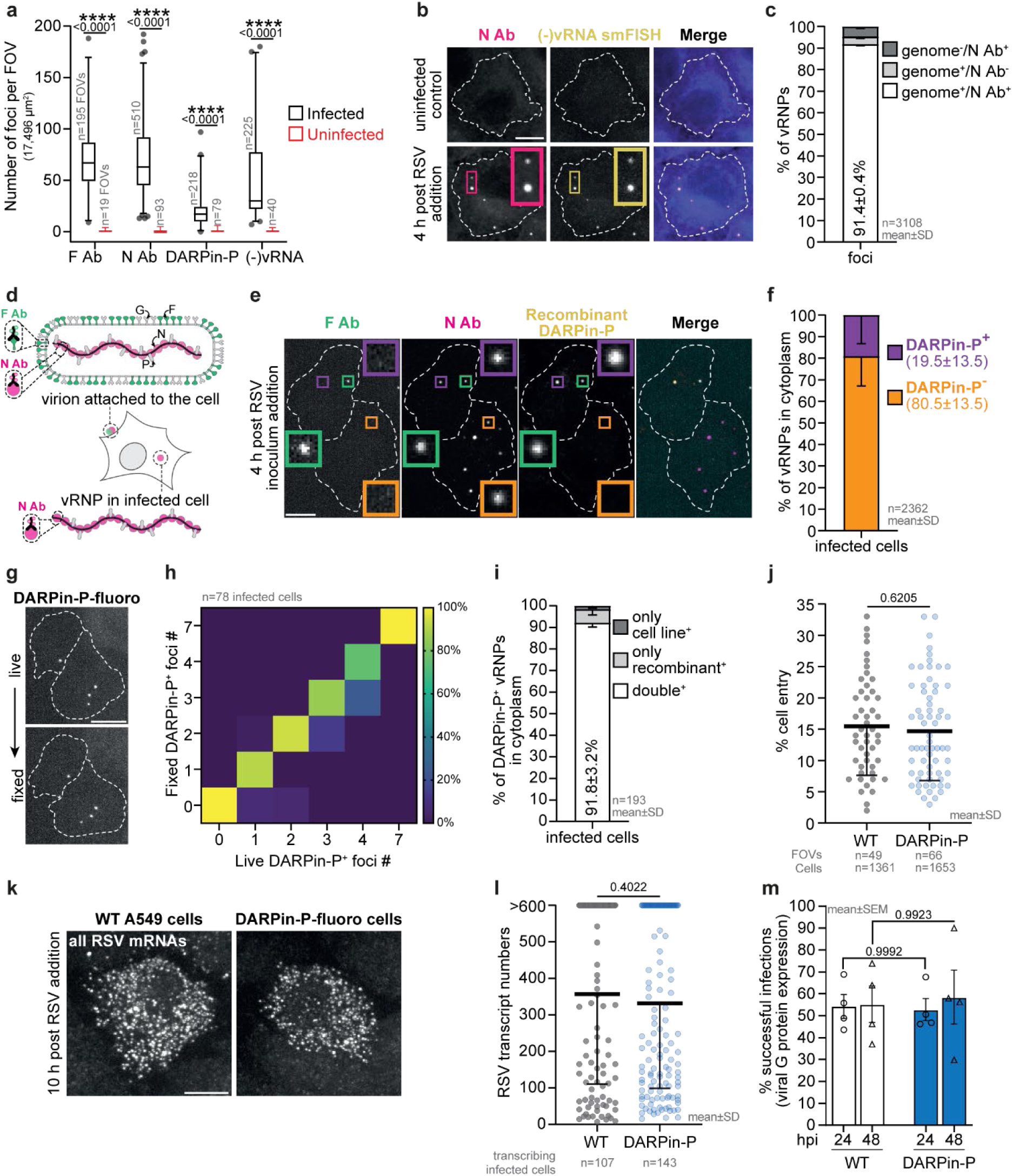
DARPin-P labels a subset of vRNPs in infected cells. **a,** Quantification of the false-positive foci detection for indicated antibodies, recombinant DARPin-P protein and RSV genome smFISH probe set used to detect RSV vRNPs and virions. **b, c,** RSV N antibody staining identifies RSV vRNPs. N antibody foci in infected cells colocalize with RSV genomic RNA ((−)vRNA). Representative images (**b**) and quantification of colocalization (**c**). **d,** Schematic highlighting dual antibody staining-based approach for distinguishing RSV vRNPs in virions (N^+^/F^+^) and in the cytoplasm of infected cells (N^+^/F^−^). **e,** Representative images of RSV infected WT A549 cells stained for RSV F, N and recombinant DARPin-P. The green insert highlights a virion attached to the cell while the orange and purple inserts highlight DARPin-P^−^ and DARPin-P^+^ vRNPs in the infected host cell cytoplasm, respectively. **f**, For cytoplasmic vRNPs (F^−^), the DARPin-P status was determined and quantified. **g, h,** The effect of paraformaldehyde fixation on DARPin-P-fluoro foci observed following RSV infection was assessed. (**g**) Representative images of the same cells imaged live and after fixation. (**h**) Heat map visualizes the frequency of the number of DARPin-P-fluoro foci observed by live imaging and imaging following fixation, in the same cells. **i**, Quantification of the colocalization between DARPin-P signal from genetically-encoded DARPin-P-fluoro and recombinant DARPin-P signal of RSV vRNPs in infected DARPin-P-fluoro cells (related to Fig. 1f). **j, k, l,** RSV infection in the DARPin-P-fluoro cell line was comparable to that in WT A549 cells. (**j**) Quantification of the fraction of infected cells at 6 h post viral inoculum addition for DARPin-P-fluoro and WT A549 cells. Representative images (**k**) and quantification (**I**) of viral mRNA transcripts (using a smFISH probe set for all RSV mRNAs) in infected DARPin-P-fluoro and WT A549 cells at 10 h post viral inoculum addition. **m,** Flow cytometry-based quantification of the percentage of infected cells expressing viral G protein (a read-out of successful infections, see **Extended Data Fig. 4h-l and Supplementary Fig. 1**) at 24 and 48 hours post viral inoculum addition for DARPin-P-fluoro and WT A549 cells. (**a, m**) Two-way ANOVA with Tukey’s multiple comparisons test used for statistical analysis. (**j, l**) Two-tailed unpaired Student’s t-test was used for statistical analysis. (**b, e, g, k**) Scale bar, 10 µm.

**Extended Data Fig 3.**
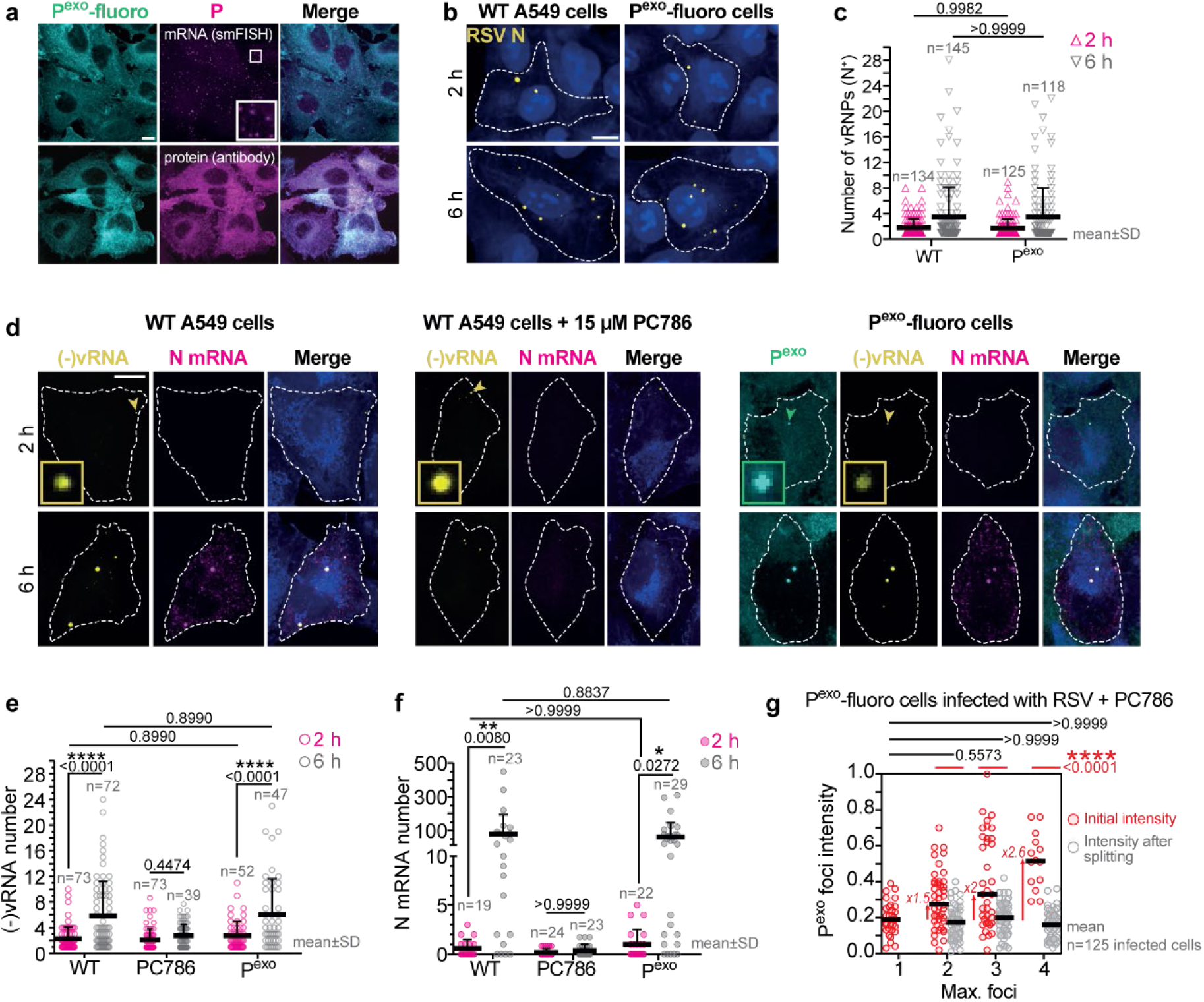
Validation of the P^exo^-fluoro cell line and RSV polymerase inhibitor, PC786. **a,** The P^exo^-fluoro cell line expresses low levels of RSV P-SunTag and fluorescently-tagged STAb. Representative images show the levels of P mRNA (via smFISH using a probe set for P) and protein (anti-RSV P antibody) in the cell line. **b, c,** RSV infection in the P^exo^-fluoro cell line was compared to that in WT A549 cells at 2 and 6 h post viral inoculum addition by antibody staining for vRNPs (anti-RSV N antibody). Representative images (**b**) and quantification of the number of vRNPs per cell (**c**) is shown. **d, e, f,** RSV infection in the P^exo^-fluoro cell line, WT A549 cells and in the presence of the RSV polymerase inhibitor, PC786, was assessed at 2 and 6 h post viral inoculum addition by smFISH staining towards the RSV genome and N transcripts. (**d**) Representative images are shown. Quantification of the (−)vRNA (**e**) and N mRNA (**f**) levels. **g,** Characterization of RSV vRNP separation after host cell entry observed for cells infected with virions containing 1-4 vRNPs. PC786 was added to all cells to ensure increases in vRNP number are caused by splitting of incoming vRNPs and not a consequence of genome replication. Foci intensities of P^exo^-fluoro for the foci before splitting (red) and individual foci intensities after reaching the maximum foci number after splitting (grey). Each data point is an individual foci. (**c, e**, **f, g**) Two-way ANOVA with Tukey’s multiple comparisons test used for statistical analysis. (**a, b, d**) Scale bar, 10 µm.

**Extended Data Fig 4.**
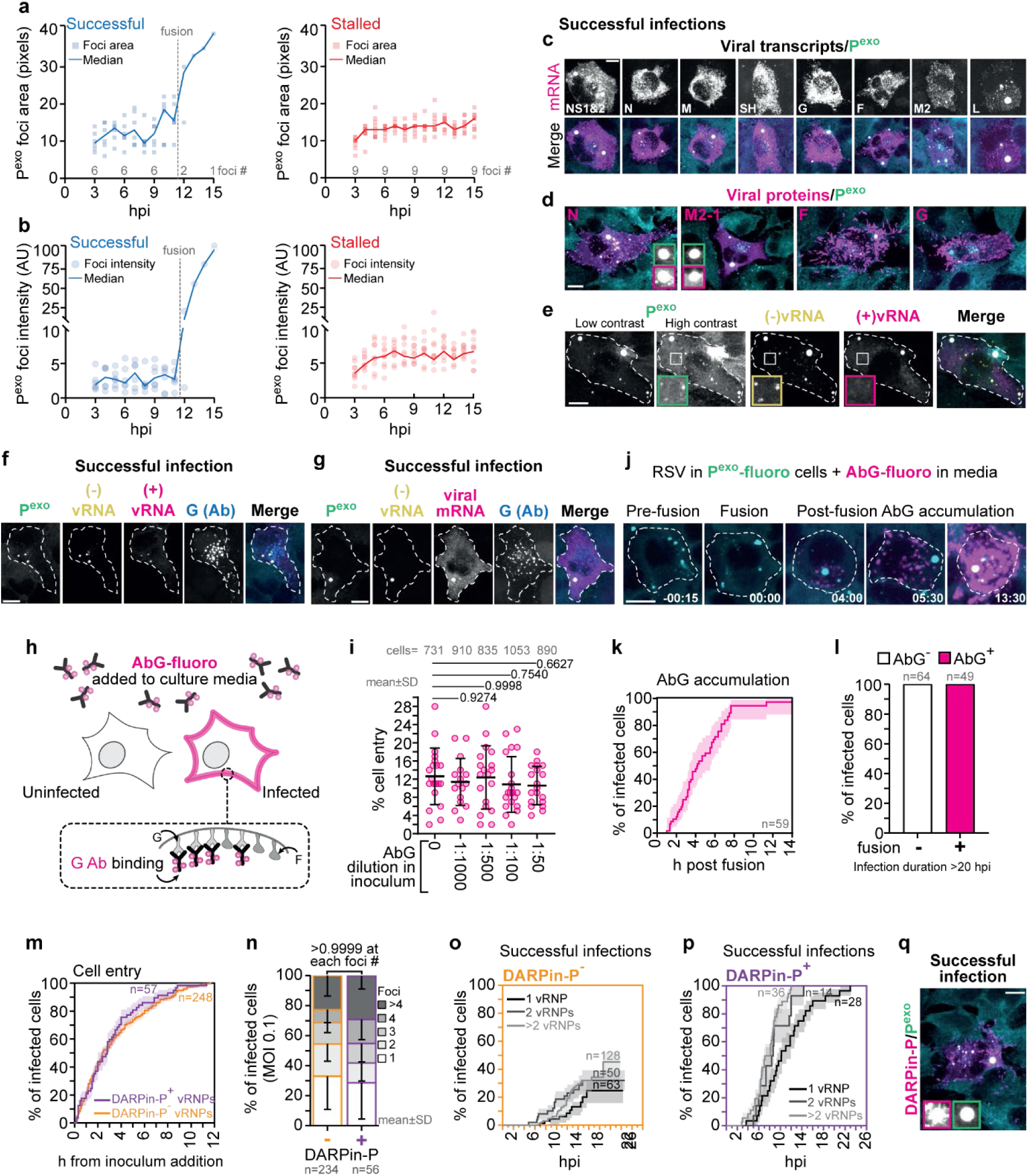
DARPin-P^+^ vRNPs give rise to successful infections. **a, b,** WT RSV infections in P^exo^-fluoro cells were further characterized. The P^exo^ foci area (**a**) and intensity (**b**) of the successful and stalled infections shown in Fig. 4a, b and **Supplementary Video 2** were quantified. Foci were assessed post splitting from 3 hpi onwards, every hour, till 15 hpi. **c, d,** Cells where a successful RSV infection was established were assessed for viral mRNA expression (**c**) and viral protein levels (**d**) in P^exo^-fluoro cells at 24 hpi. **e,** Following IB formation in RSV infected P^exo^-fluoro cells, there is a rapid increase in foci. These are vRNP progeny that stain positive for RSV genomic RNA by smFISH. **f, g,** RSV infected cells that were positive for extracellular viral G protein were assessed for their viral RNA composition by smFISH. These cells always displayed viral antigenomes (**f**) as well as high levels of viral mRNA transcripts (RSV whole transcriptome smFISH probe set) (**g**). **h, i, j, k, l,** Cell culture supplemented fluorescently-conjugated RSV G antibody (AbG-fluoro) identifies successful RSV infections (as defined by observing IBs and vRNP replication). (**h**) Schematic of the AbG-fluoro labeling approach. (**i**) P^exo^-fluoro cells were infected with viral inoculum supplemented with increasing concentrations of AbG-fluoro and infection assessed at 6 hpi. Quantification shows no inhibitory effect on viral entry at concentrations up to 1:50 of the antibody compared to no antibody control. Each data point represents an imaged FOV. (**j**) Representative images of extracellular AbG-fluoro accumulation on infected cells relative to the moment of vRNP fusion. (**k**) The timing of AbG-fluoro labeling relative to vRNP fusion is displayed by a Kaplan-Meier graph. Only infections with >1 vRNP were included to analyze vRNP fusion. (**l**) For infections with >1 vRNP, AbG-fluoro positivity is exclusively observed in infections where vRNPs have fused. All cells were imaged for at least 20 hpi. **m, n,** The potential variations in infection by virions carrying DARPin-P^+^ and DARPin-P^−^ vRNPs were assessed. (**m**) Kaplan-Meier curves showing the time from inoculum addition to vRNP entry for both DARPin-P^+^ and DARPin-P^−^ virions. (**n**) The number of vRNPs per infected cell for DARPin-P^+^ and DARPin-P^−^ virions was quantified. Comparing DARPin-P^+^ and DARPin-P^−^ virions show no significant difference across all foci numbers. **o, p,** Infection success for DARPin-P^−^ (**o**) and DARPin-P^+^ (**p**) infections were assessed in relation to the number of vRNPs and displayed as Kaplan-Meier graphs. **q,** DARPin-P status of successfully infected cells was assessed. All IBs in these infections are DARPin-P positive. (**k, m, o, p**) Lines and shaded areas indicate mean and SE, respectively. (**i**) Ordinary one-way ANOVA with Dunnett’s multiple comparisons test used for statistical analysis. (**n**) Two-way ANOVA with Tukey’s multiple comparisons test used for statistical analysis. (**c, d, e, f, g, j, q**) Scale bar, 10 µm. (**j**) Time, h: min.

**Extended Data Fig 5.**
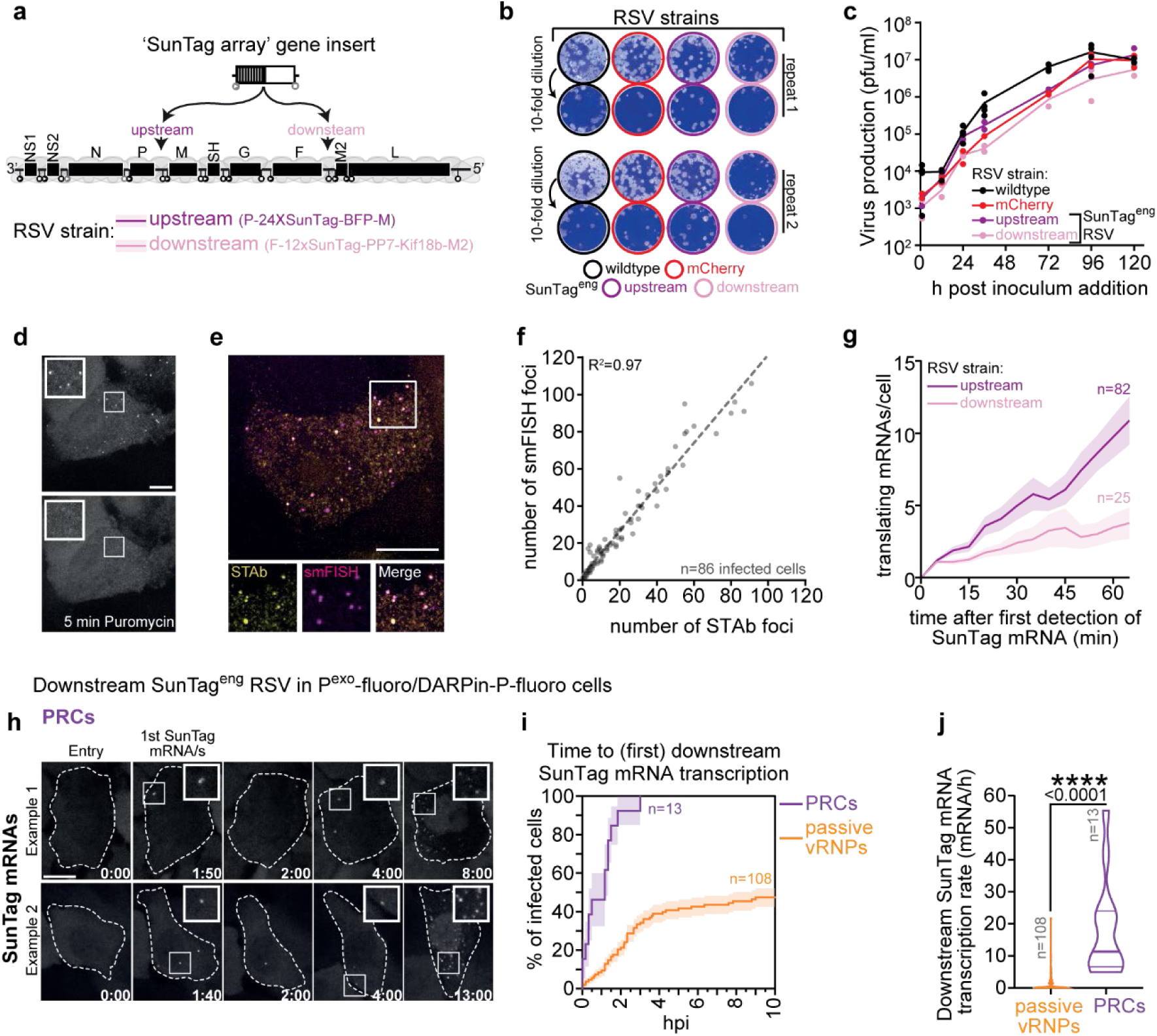
Validation of the SunTag^eng^ RSV strains that allow real-time visualization of viral mRNA expression. **a,** Schematic illustrating the upstream and downstream insertion locations of the ‘additional SunTag genes’ in the two SunTag^eng^ RSV strains. **b, c,** Viral growth analysis based on plaque assays were carried out to determine the fitness of the engineered RSV strains compared to the WT. (**b**) Representative images of observed plaques 6 days post viral inoculation. For each viral strain, 2 repeats, each with 2 inoculum dilutions are shown. (**c**) Quantification of growth assay. Line connects mean of experimental repeats (dots). **d,** Representative images of time-lapse movies of cells expressing STAb-fluoro after infection with upstream SunTag^eng^ RSV prior to (top panel) and following (bottom panel) puromycin administration. **e, f,** STAb-fluoro foci observed with the SunTag^eng^ RSV strains were assessed by smFISH. (**e**) Representative images of smFISH for SunTag mRNA foci and STAb-fluoro foci in the same infected cell for the upstream strain is shown. (**f**) Scatter dot plot quantifies the number of SunTag mRNAs (via smFISH) and translation sites (via STAb) in the same cells. Each dot indicates a single cell and dashed line indicates linear correlation with Pearson R^2^ (top left). **g,** The number of translating SunTag mRNAs over time was quantified for both the upstream and downstream SunTag^eng^ strains. Overall, both strains show an increase in the number of translating SunTag mRNAs over time. Mean (line) and SE (shaded area) are shown. **h, i, j,** Downstream SunTag^eng^ RSV viral mRNA expression dynamics were assessed for infections originating from passive vRNPs and PRCs in cell lines labelling all vRNPs (P^exo^-fluoro) and DARPin-P^+^ vRNPs (DARPin-P-fluoro). (**h**) Two representative image series from time-lapse movies of SunTag mRNAs in cells infected with a PRC are shown. (**i**) Kaplan-Meier graphs depict the start of viral transcription for infections with passive vRNPs and PRCs. Mean (line) and SE (shaded area) are shown. (**j**) The transcription rate was calculated and plotted as violin plots with the median and quartiles shown (horizontal lines). Two-tailed unpaired Student’s t-test was used for statistical analysis. (**d, e, h**) Scale bar, 10 µm. (**h**) Time, h: min.

**Extended Data Fig 6.**
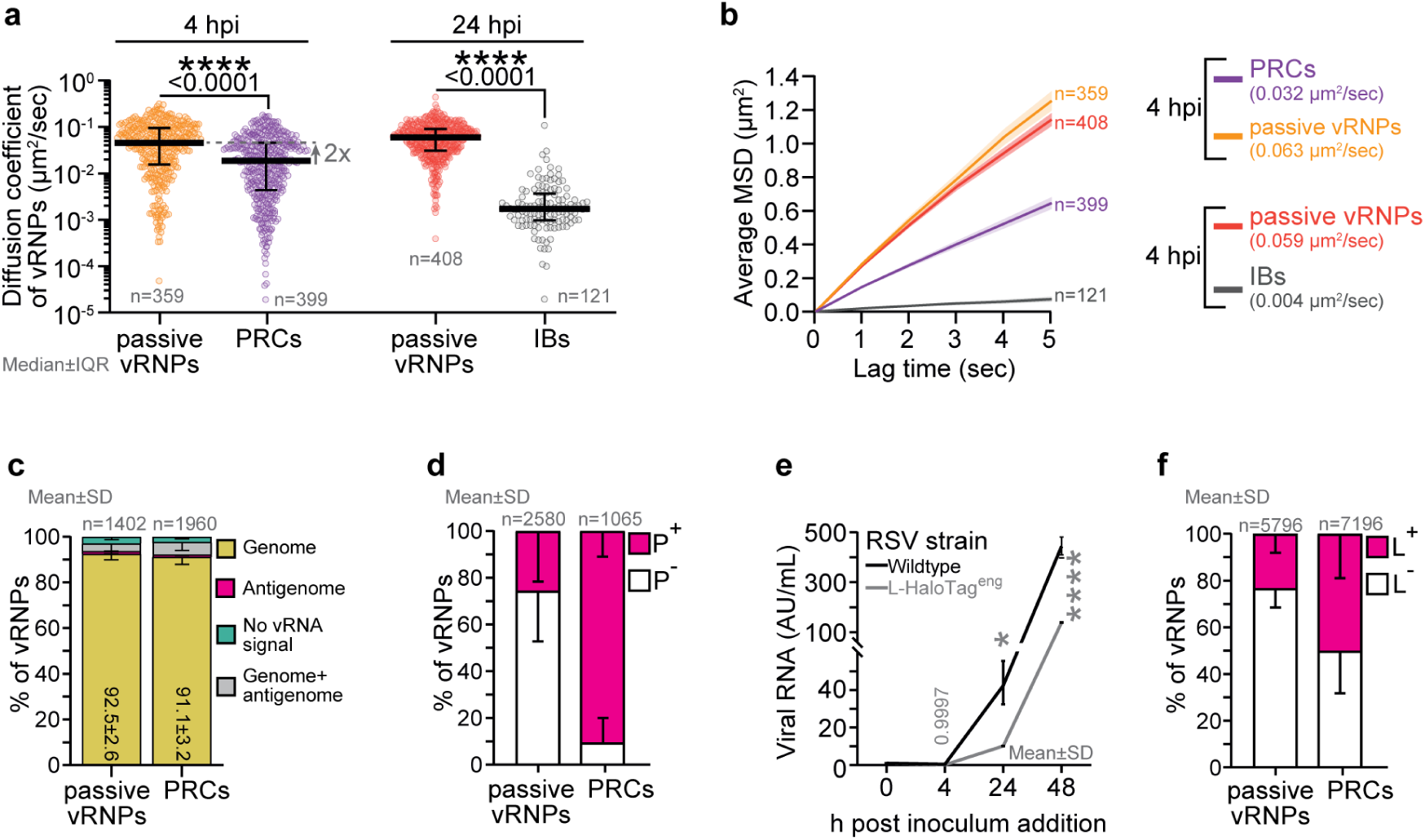
Characterization of vRNP diffusion and protein composition. **a, b,** The mobility of infecting vRNPs (RSV and emetine added for 4 h), IBs in successful infections and vRNPs in stalled infections (RSV added for 24 h) was assessed in the P^exo^-fluoro/DARPin-P-fluoro cell line. (**a**) The diffusion coefficient of individual foci is plotted. (**b**) The mobility data of foci at 4 and 24 hpi is visualized as the average MSD of all vRNPs over a 5 sec time period. **c,** The frequency of the different vRNA species in infecting vRNPs is quantified (related to Fig. 6a). **d,** Frequency of passive vRNPs and PRC foci that stains positive for P antibody stain is quantified (related to Fig. 6e, f). **e,** Viral growth analysis based on qPCR assays was carried out to determine the fitness of the engineered L-HaloTag^eng^ RSV strain compared to the WT strain. **f,** Frequency of passive vRNPs and PRC foci that are positive for L-HaloTag signal is quantified (related to Fig. 6h, i). (**b**) Mean (line) and SE (shaded area) are shown. (**a**) Two-tailed unpaired Student’s t-test was used for statistical analysis. (**e**) Two-way ANOVA with Dunnett’s multiple comparisons test was used for statistical analysis.

**Extended Data Fig 7.**
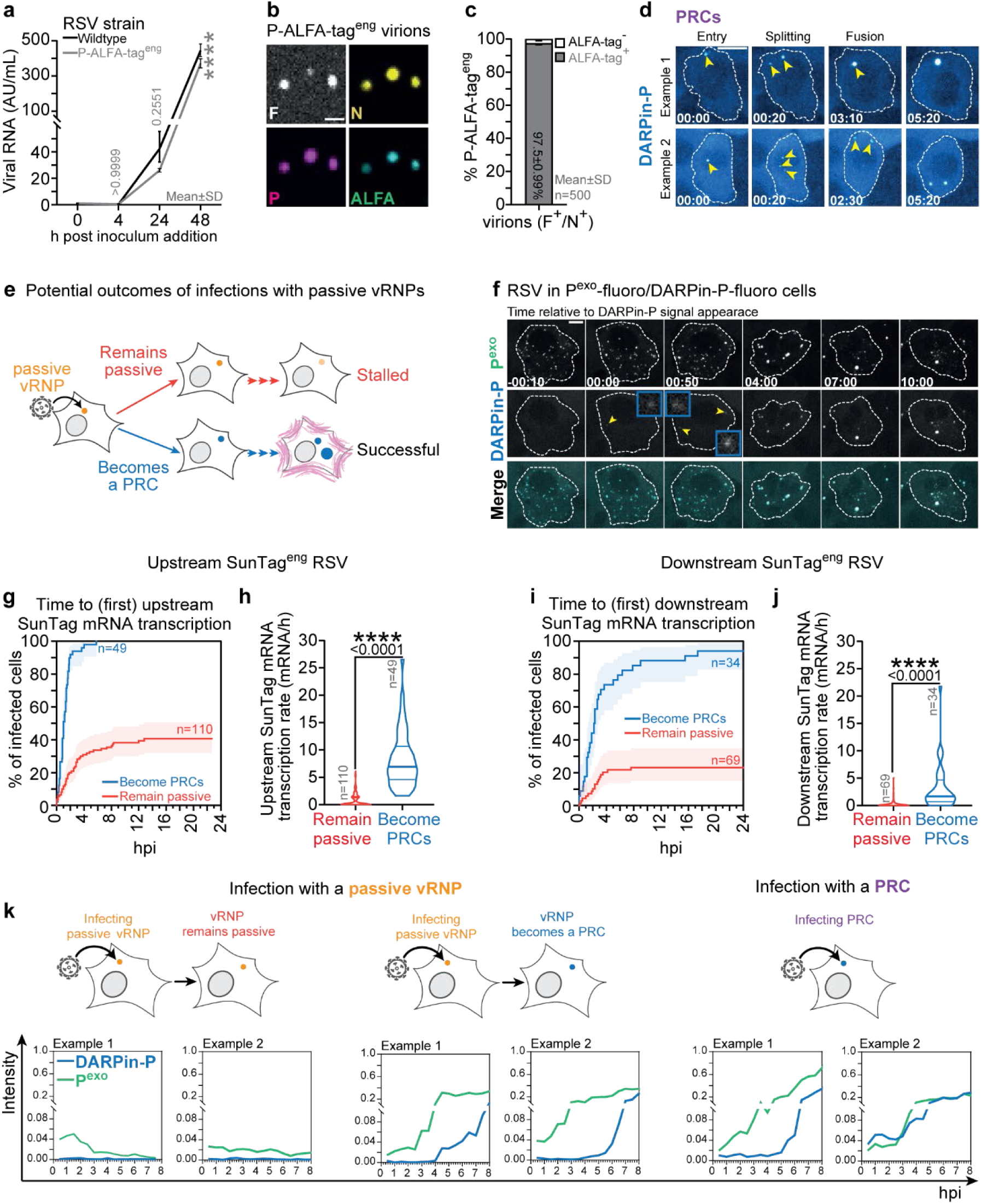
Characterizing infections where vRNPs become PRCs post entry. **a, b, c,** The P-ALFA-Tag^eng^ RSV strain encodes for a single ALFA-tag in the P gene. This strain enables fluorescent-labelling of endogenous viral P protein (expressed from the viral genome) upon infection of cells expressing Nb ALFA-fluoro. (**a**) Viral growth analysis based on qPCR assays were carried out to determine the fitness of the engineered P-ALFA-Tag^eng^ RSV strain compared to the WT strain. Representative images (**b**) and quantification of the fraction of virions that are positive for the ALFA-tag (**c**) in P-ALFA-Tag^eng^ RSV virion population. ALFA-tag was detected using an anti-ALFA-tag antibody. **d,** Two representative image series from time-lapse movies showing infections with PRCs, where the vRNPs initially split, then subsequently fuse and increase in intensity. **e, f,** Infections with passive vRNPs were further characterized. (**e**) Schematic highlighting the potential outcomes of passive vRNP infections. (**f**) Representative image series from a time-lapse movie showing an infection with multiple passive vRNPs where one vRNP becomes a PRC post cell entry. Yellow arrowheads indicate first appearance of DARPin-P positivity. **g, h, i, j,** Analysis of the transcriptional activity of passive vRNP infections that either become PRCs or remain passive. Kaplan-Meier graphs depicting the start of viral transcription (**g**) and violin plots showing the transcription rate (**h**) for the upstream SunTag^eng^ RSV strain. Kaplan-Meier graphs depicting the start of viral transcription (**i**) and violin plots showing the transcription rate (**j**) for the downstream SunTag^eng^ RSV strain. **k,** Example intensity-time traces of vRNP P^exo^-fluoro and DARPin-P-fluoro foci intensities are shown for the following 3 infection scenarios: (1) infections arising from passive^−^ vRNPs that remain passive, (2) infections from passive vRNPs that becomes a PRC during infection and, (3) infections with a PRC. (**g, i**) Mean (line) and SE (shaded area) are shown. (**h, j**) The median and quartiles are shown (horizontal lines) (**a**) Two-way ANOVA with Dunnett’s multiple comparisons test was used for statistical analysis. (**h, j**) Two-tailed unpaired Student’s t-test was used for statistical analysis. (**b**) Scale bars, 2 µm. (**d, f**) Scale bars, 10 µm and time, h: min.

## Supplementary information

**Supplementary Table 1.** Number of experimental repeats, details of cell lines, antibodies and smFISH probe sets, oligonucleotides and construct sequences related to methods.

**Supplementary Video 1. Real-time observations of vRNP release into host cell cytoplasm upon infection by RSV virions with different numbers of vRNPs.**

Maximum intensity projections of P^exo^-fluoro cells infected with WT RSV in the presence of 15 µM PC786 (RSV polymerase inhibitor) to inhibit viral replication. Movie highlights 6 individual cells of interest infected with virions carrying different numbers of vRNPs. The max vRNP number for each infection is stated. Images were acquired every 5 min and the first image where viral foci are observed is set to t=0. The individual movies stop once the maximum foci number is reached and the cell infected with a virion carrying 1 foci stops at 3 h. Representative images from the time-lapse movie of the cells infected with virions carrying 2, 4 and 8 vRNPs are show in **Fig. 2g**.

**Supplementary Video 2. Time-lapse imaging of RSV infection establishment with single vRNP resolution.**

Maximum intensity projection of P^exo^-fluoro cells infected with WT RSV. Movie highlights one cell undergoing a successful infection with vRNP fusion and vRNP replication (left) and one with a stalled infection that fails to undergo viral replication (right). Frame freezes with indicative arrows have been used to enhance clarity and highlight key events of the viral life cycle. Images were acquired every 10 min and the first image where viral foci are observed is set to t=0. Representative images from the time-lapse movie are show in in **Fig. 3a**, foci number traces are show in **Fig. 3b** and foci area and intensity are quantified in **Extended Data Fig. 4a** and b, respectively.

**Supplementary Video 3. Live observation of DARPin-P^−^ and DARPin-P^+^ vRNP-driven RSV infections.**

Maximum intensity projection of cells stably expressing P^exo^-fluoro and DARPin-P-fluoro infected with an inoculum containing WT RSV and AbG-fluoro. Movie highlights two individual cells of interest, one infected with a DARPin-P^−^ vRNP (left) and the other infected with a DARPin-P^+^ vRNP (right). The individual channels (greyscale) and merge are shown. Images were acquired every 20 min and the first image where viral foci are observed is set to t=0. The infecting vRNP is highlighted in the P^exo^-fluoro channel for both DARPin-P negative and positive vRNPs and the DARPIn-P-fluoro channel for only the DARPin-P positive vRNP. Representative images from the time-lapse movie are show in **Fig. 3i**.

**Supplementary Video 4. Real-time visualization of viral mRNA transcription using the upstream and downstream SunTag^eng^ RSV strain.**

Maximum intensity projection of P^exo^-fluoro/DARPin-H6-fluoro/STAb-fluoro cells infected with the upstream (top row) and downstream (bottom row) SunTag^eng^ RSV strains. For each viral strain an infection with a passive vRNP (left) and PRC (right) are highlighted. The video shows only the STAb-fluoro channel. Images were acquired every 10 min and the first image where vRNPs are observed is set to t=0. Arrows highlight the first frame where STAb foci are observed. Representative images from the upstream strain time-lapse movies are show in **Fig. 4b, c**.

**Supplementary Video 5. Live visualization of a passive vRNP becoming a PRC during the infection process.**

Maximum intensity projection of cells stably expressing P^exo^-fluoro and DARPin-P-fluoro infected with an inoculum containing WT RSV and AbG-fluoro. Video highlights a cell infected by a virion carrying a single passive vRNP that goes on to become a PRC. Images were acquired every 10 min and the first image where a viral vRNP is observed is set to t=0. The vRNP at infection and upon acquiring DARPin-P positivity is highlighted with arrows.

**Supplementary Fig 1.**
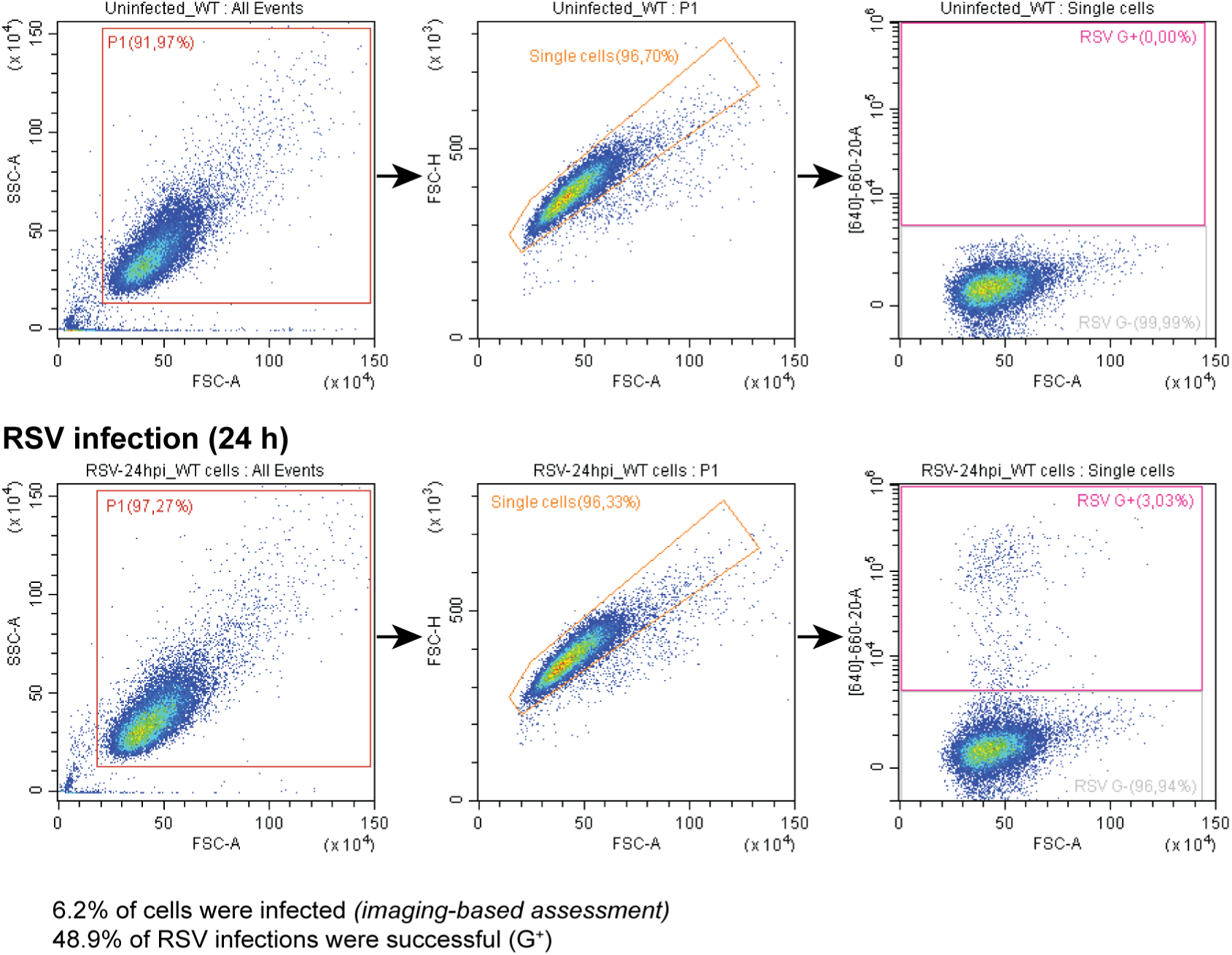
Flow cytometry gating strategy for successful infections. Successful RSV infections result in extracellular expression of viral G protein. These cells were identified by flow cytometry at the time points of interest and related to the total number of infected cells (established by imaging) to calculate the frequency of infection success. Uninfected control cells were used as negative control to aid in selecting G positive cells in the RSV infected sample. Cells were separated from debris by analyzing forward scatter versus side scatter. Subsequently, single cells were identified by plotting forward scatter-area against forward scatter-height. Uninfected control cells were used to identify cells negative for RSV G and then infected cells were analyzed.

## References

1 Rincheval, V. et al. Functional organization of cytoplasmic inclusion bodies in cells infected by respiratory syncytial virus. Nature communications 8, 563 (2017).

2 Lahaye, X. et al. Functional characterization of Negri bodies (NBs) in rabies virus-infected cells: Evidence that NBs are sites of viral transcription and replication. Journal of virology 83, 7948–7958 (2009).

3 Hoenen, T. et al. Inclusion bodies are a site of ebolavirus replication. Journal of virology 86, 11779–11788 (2012).

4 Zhou, Y., Su, J. M., Samuel, C. E. & Ma, D. Measles virus forms inclusion bodies with properties of liquid organelles. Journal of virology 93, 10.1128/jvi.00948-00919 (2019).

5 Mitrea, D. M., Mittasch, M., Gomes, B. F., Klein, I. A. & Murcko, M. A. Modulating biomolecular condensates: a novel approach to drug discovery. Nature Reviews Drug Discovery 21, 841–862 (2022).

6 Bakker, S. E. et al. The respiratory syncytial virus nucleoprotein–RNA complex forms a left-handed helical nucleocapsid. Journal of General Virology 94, 1734–1738 (2013).

7 Tawar, R. G. et al. Crystal structure of a nucleocapsid-like nucleoprotein-RNA complex of respiratory syncytial virus. Science 326, 1279–1283 (2009).

8 Šantak, M. & Matić, Z. The role of nucleoprotein in immunity to human negative-stranded RNA viruses—not just another brick in the viral nucleocapsid. Viruses 14, 521 (2022).

9 Yu, Q., Hardy, R. W. & Wertz, G. W. Functional cDNA clones of the human respiratory syncytial (RS) virus N, P, and L proteins support replication of RS virus genomic RNA analogs and define minimal trans-acting requirements for RNA replication. Journal of Virology 69, 2412–2419 (1995).

10 Collins, P. L. et al. Production of infectious human respiratory syncytial virus from cloned cDNA confirms an essential role for the transcription elongation factor from the 5’proximal open reading frame of the M2 mRNA in gene expression and provides a capability for vaccine development. Proceedings of the National Academy of Sciences 92, 11563–11567 (1995).

11 García, J., García-Barreno, B., Vivo, A. & Melero, J. A. Cytoplasmic inclusions of respiratory syncytial virus-infected cells: formation of inclusion bodies in transfected cells that coexpress the nucleoprotein, the phosphoprotein, and the 22K protein. Virology 195, 243–247 (1993).

12 Galloux, M. et al. Minimal elements required for the formation of respiratory syncytial virus cytoplasmic inclusion bodies in vivo and in vitro. MBio 11, 10.1128/mbio.01202-01220 (2020).

13 Alberti, S. Phase separation in biology. Current Biology 27, R1097–R1102 (2017).

14 Rabouw, H. H. et al. Mapping the complete influenza A virus infection cycle through single vRNP imaging. bioRxiv, 2025.2001.2020.633851, doi:10.1101/2025.01.20.633851 (2025).

15 Boersma, S. et al. Translation and replication dynamics of single RNA viruses. Cell 183, 1930–1945. e1923 (2020).

16 Rameix-Welti, M.-A. et al. Visualizing the replication of respiratory syncytial virus in cells and in living mice. Nature communications 5, 1–10 (2014).

17 Plückthun, A. Designed ankyrin repeat proteins (DARPins): binding proteins for research, diagnostics, and therapy. Annual review of pharmacology and toxicology 55, 489–511 (2015).

18 Galloux, M. et al. Characterization of a viral phosphoprotein binding site on the surface of the respiratory syncytial nucleoprotein. Journal of virology 86, 8375–8387 (2012).

19 Tanenbaum, M. E., Gilbert, L. A., Qi, L. S., Weissman, J. S. & Vale, R. D. A protein-tagging system for signal amplification in gene expression and fluorescence imaging. Cell 159, 635–646 (2014).

20 Wörn, A. et al. Correlation between in vitro stability and in vivo performance of anti-GCN4 intrabodies as cytoplasmic inhibitors. Journal of Biological Chemistry 275, 2795–2803 (2000).

21 Coates, M. et al. Preclinical characterization of PC786, an inhaled small-molecule respiratory syncytial virus L protein polymerase inhibitor. Antimicrobial Agents and Chemotherapy 61, 10.1128/aac.00737-00717 (2017).

22 Kiss, G. et al. Structural analysis of respiratory syncytial virus reveals the position of M2-1 between the matrix protein and the ribonucleoprotein complex. Journal of virology 88, 7602–7617 (2014).

23 Loney, C., Mottet-Osman, G., Roux, L. & Bhella, D. Paramyxovirus ultrastructure and genome packaging: cryo-electron tomography of sendai virus. Journal of virology 83, 8191–8197 (2009).

24 Liljeroos, L., Krzyzaniak, M. A., Helenius, A. & Butcher, S. J. Architecture of respiratory syncytial virus revealed by electron cryotomography. Proceedings of the National Academy of Sciences 110, 11133–11138 (2013).

25 Donovan-Banfield, I. a., et al. Direct RNA sequencing of respiratory syncytial virus infected human cells generates a detailed overview of RSV polycistronic mRNA and transcript abundance. Plos one 17, e0276697 (2022).

26 Götzke, H. et al. The ALFA-tag is a highly versatile tool for nanobody-based bioscience applications. Nature communications 10, 4403 (2019).

27 Gonnin, L. et al. Structural landscape of the respiratory syncytial virus nucleocapsids. Nature Communications 14, 5732 (2023).

28 Los, G. V. et al. HaloTag: a novel protein labeling technology for cell imaging and protein analysis. ACS chemical biology 3, 373–382 (2008).

29 Gilman, M. S. et al. Structure of the respiratory syncytial virus polymerase complex. Cell 179, 193–204. e114 (2019).

30 Norrby, E., Marusyk, H. & Örvell, C. Morphogenesis of respiratory syncytial virus in a green monkey kidney cell line (Vero). Journal of virology 6, 237–242 (1970).

31 García-Barreno, B., Delgado, T. & Melero, J. A. Identification of protein regions involved in the interaction of human respiratory syncytial virus phosphoprotein and nucleoprotein: significance for nucleocapsid assembly and formation of cytoplasmic inclusions. Journal of virology 70, 801–808 (1996).

32 Carromeu, C., Simabuco, F. M., Tamura, R. E., Farinha Arcieri, L. E. & Ventura, A. Intracellular localization of human respiratory syncytial virus L protein. Archives of virology 152, 2259–2263 (2007).

33 Lindquist, M. E., Lifland, A. W., Utley, T. J., Santangelo, P. J. & Crowe Jr, J. E. Respiratory syncytial virus induces host RNA stress granules to facilitate viral replication. Journal of virology 84, 12274–12284 (2010).

34 Santangelo, P. J. & Bao, G. Dynamics of filamentous viral RNPs prior to egress. Nucleic acids research 35, 3602–3611 (2007).

35 Lifland, A. W. et al. Human respiratory syncytial virus nucleoprotein and inclusion bodies antagonize the innate immune response mediated by MDA5 and MAVS. Journal of virology 86, 8245–8258 (2012).

36 Cardone, C. et al. A structural and dynamic analysis of the partially disordered polymerase-binding domain in RSV phosphoprotein. Biomolecules 11, 1225 (2021).

37 Simabuco, F. M. et al. Structural analysis of human respiratory syncytial virus p protein: identification of intrinsically disordered domains. Brazilian Journal of Microbiology 42, 340–345 (2011).

38 Brocca, S., Grandori, R., Longhi, S. & Uversky, V. Liquid–liquid phase separation by intrinsically disordered protein regions of viruses: Roles in viral life cycle and control of virus–host interactions. International Journal of Molecular Sciences 21, 9045 (2020).

39 Su, J. M., Wilson, M. Z., Samuel, C. E. & Ma, D. Formation and function of liquid-like viral factories in negative-sense single-stranded RNA virus infections. Viruses 13, 126 (2021).

40 Liang, B. Structures of the mononegavirales polymerases. Journal of Virology 94, 10.1128/jvi.00175-00120 (2020).

41 Buchholz, U. J., Finke, S. & Conzelmann, K.-K. Generation of bovine respiratory syncytial virus (BRSV) from cDNA: BRSV NS2 is not essential for virus replication in tissue culture, and the human RSV leader region acts as a functional BRSV genome promoter. Journal of virology 73, 251–259 (1999).

42 Pereira, N. et al. New insights into structural disorder in human respiratory syncytial virus phosphoprotein and implications for binding of protein partners. Journal of Biological Chemistry 292, 2120–2131 (2017).

43 Dreier, B. & Plückthun, A. in Ribosome Display and Related Technologies: Methods and Protocols 261–286 (Springer, 2011).

44 Binz, H. K., Stumpp, M. T., Forrer, P., Amstutz, P. & Plückthun, A. Designing repeat proteins: well-expressed, soluble and stable proteins from combinatorial libraries of consensus ankyrin repeat proteins. Journal of molecular biology 332, 489–503 (2003).

45 Kramer, M. A., Wetzel, S. K., Plückthun, A., Mittl, P. R. & Grütter, M. G. Structural determinants for improved stability of designed ankyrin repeat proteins with a redesigned C-capping module. Journal of molecular biology 404, 381–391 (2010).

46 Brauchle, M. et al. Protein interference applications in cellular and developmental biology using DARPins that recognize GFP and mCherry. Biology open 3, 1252–1261 (2014).

47 Schilling, J., Schöppe, J. & Plückthun, A. From DARPins to LoopDARPins: novel LoopDARPin design allows the selection of low picomolar binders in a single round of ribosome display. Journal of molecular biology 426, 691–721 (2014).

48 Zahnd, C., Sarkar, C. A. & Plückthun, A. Computational analysis of off-rate selection experiments to optimize affinity maturation by directed evolution. Protein Engineering, Design & Selection 23, 175–184 (2010).

49 Campbell, B. F., Dittmann, A., Dreier, B., Plückthun, A. & Tyagarajan, S. K. A DARPin-based molecular toolset to probe gephyrin and inhibitory synapse biology. Elife 11, e80895 (2022).

50 Simon, M., Zangemeister-Wittke, U. & Plückthun, A. Facile double-functionalization of designed ankyrin repeat proteins using click and thiol chemistries. Bioconjugate chemistry 23, 279–286 (2012).

51 Skinner, S. P. et al. CcpNmr AnalysisAssign: a flexible platform for integrated NMR analysis. Journal of biomolecular NMR 66, 111–124 (2016).

52 Gonnin, L. et al. Importance of RNA length for in vitro encapsidation by the nucleoprotein of human respiratory syncytial virus. Journal of Biological Chemistry 298 (2022).

53 Risso-Ballester, J. et al. A condensate-hardening drug blocks RSV replication in vivo. Nature 595, 596–599 (2021).

54 Fix, J., Galloux, M., Blondot, M.-L. & Eléouët, J.-F. The insertion of fluorescent proteins in a variable region of respiratory syncytial virus L polymerase results in fluorescent and functional enzymes but with reduced activities. The open virology journal 5, 103 (2011).

55 Blanchard, E. L. et al. Polymerase-tagged respiratory syncytial virus reveals a dynamic rearrangement of the ribonucleocapsid complex during infection. PLoS pathogens 16, e1008987 (2020).

56 Thi Nhu Thao, T., et al. Rapid reconstruction of SARS-CoV-2 using a synthetic genomics platform. Nature 582, 561–565 (2020).

57 Gibson, D. G. et al. Creation of a bacterial cell controlled by a chemically synthesized genome. science 329, 52–56 (2010).

58 Kouprina, N., Noskov, V. N. & Larionov, V. Selective isolation of large segments from individual microbial genomes and environmental DNA samples using transformation-associated recombination cloning in yeast. Nature protocols 15, 734–749 (2020).

59 Bouillier, C. et al. Generation, amplification, and titration of recombinant respiratory syncytial viruses. JoVE (Journal of Visualized Experiments), e59218 (2019).

60 Grimm, J. B. et al. A general method to improve fluorophores using deuterated auxochromes. Jacs Au 1, 690–696 (2021).

61 Gaspar, I., Wippich, F. & Ephrussi, A. Terminal deoxynucleotidyl transferase mediated production of labeled probes for single-molecule FISH or RNA capture. Bio-protocol 8, e2750–e2750 (2018).

62 Lyubimova, A. et al. Single-molecule mRNA detection and counting in mammalian tissue. Nature protocols 8, 1743–1758 (2013).

63 Khuperkar, D. et al. Quantification of mRNA translation in live cells using single-molecule imaging. Nature Protocols 15, 1371–1398 (2020).

64 Model, M. A. & Burkhardt, J. K. A standard for calibration and shading correction of a fluorescence microscope. Cytometry: The Journal of the International Society for Analytical Cytology 44, 309–316 (2001).

65 Van der Walt, S. et al. scikit-image: image processing in Python. PeerJ 2, e453 (2014).

66 Fukai, Y. T. & Kawaguchi, K. LapTrack: linear assignment particle tracking with tunable metrics. Bioinformatics 39, btac799 (2023).

67 Ahlers, J. et al. napari: a multi-dimensional image viewer for Python. Zenodo, 1–2 (2023).

68 Ritchie, M. E. et al. limma powers differential expression analyses for RNA-sequencing and microarray studies. Nucleic acids research 43, e47–e47 (2015).

69 Tran, T.-L. et al. The nine C-terminal amino acids of the respiratory syncytial virus protein P are necessary and sufficient for binding to ribonucleoprotein complexes in which six ribonucleotides are contacted per N protein protomer. Journal of general virology 88, 196–206 (2007).

70 Abramson, J. et al. Accurate structure prediction of biomolecular interactions with AlphaFold 3. Nature, 1–3 (2024).

